# Geographic population structure and distinct population dynamics of globally abundant freshwater bacteria

**DOI:** 10.1101/2023.07.13.548520

**Authors:** M. Hoetzinger, M.W. Hahn, L.Y. Andersson, N. Buckley, C. Ramsin, M. Buck, J.K. Nuy, S.L. Garcia, F. Puente-Sánchez, S. Bertilsson

**Affiliations:** Department of Aquatic Sciences and Assessment, Swedish University of Agricultural Sciences, Uppsala, Sweden; Faculty of Chemistry, Biotechnology and Food Science, Norwegian University of Life Sciences, Ås, Norway; Research Department for Limnology, University of Innsbruck, Mondsee, Austria; Department of Ecology, Environment, and Plant Sciences, Science for Life Laboratory, Stockholm University, Stockholm, Sweden; Centre for Water and Environmental Research, University of Duisburg-Essen, Essen, Germany

## Abstract

Geographic separation is a principal factor for structuring populations of macroorganisms, with important consequences for evolution, by means of processes such as allopatric speciation. For free-living prokaryotes, implications of geographic separation on their evolution are more unclear. The limited phylogenetic resolution of commonly used markers such as 16S rRNA gene sequences have since long impeded prokaryotic population genetics. However, the vast amount of metagenome sequencing data generated during the last decades from various habitats around the world, now provides an excellent opportunity for such investigations. Here we exploited publicly available and new freshwater metagenomes in combination with genomes of abundant freshwater bacteria to study the impact of geographic separation on population structure. We focused on species that were detected across broad geographic ranges at high enough sequence coverage for meaningful population genomic analyses, i.e. members of the predominant freshwater taxa acI, LD12, *Polynucleobacter* and *Ca*. Methylopumilus. Population differentiation increased significantly with spatial distance in all species, but notable dispersal barriers (e.g. oceanic) were not apparent. Yet, the different species showed contrasting rates of geographic divergence and strikingly different population dynamics in time series within individual lakes. While certain populations hardly diverged over several years, others displayed high divergence after merely a few months, similar in scale to populations separated by thousands of kilometers. We speculate that populations with higher strain diversity evolve more monotonously, while low strain diversity enables more drastic clonal expansion of genotypes which will be reflected in strong but transient differentiation between temporally or spatially adjacent populations.

## Introduction

The importance of geographic separation for evolution has long been recognized: “…barriers of any kind, or obstacles to free migration, are related in a close and important manner to the differences between the productions of various regions.” (Darwin, 1859). However, in comparison to most macroorganisms, bacteria are exceedingly mobile due to their minuscule size, which facilitates long-range dispersal, e.g. via clouds (Griffin, 2007; Smith *et al*., 2012; DeLeon-Rodriguez *et al*., 2013; Sarmiento-Vizcaíno *et al*., 2018), ocean currents (Müller *et al*., 2014; Hellweger *et al*., 2014; Whittaker and Rynearson, 2017) or by hitchhiking with larger migratory organisms (Reed *et al*., 2003; Grossart *et al*., 2010; Comas *et al*., 2013). Examples of entire orders being restricted to certain continents, as is the case for several mammals (Smith, 1983), are thus not to be expected. Still, geographic endemism occurs in bacteria, albeit at higher phylogenetic resolution. A recent study relating average nucleotide identity (ANI) between publicly available genomes to geographic distance between their sites of origin provided valuable insights into the phylogenetic scales at which prokaryotes from different environments display endemism (Louca, 2021). On one end of the spectrum, prokaryotes from the subsurface showed the highest level of endemism. No clades above the commonly used species delineation threshold of 95% ANI (Konstantinidis *et al*., 2006; Jain *et al*., 2018) were found on opposite hemispheres of the earth. On the other hand, for marine environments, clades were not restricted to a single hemisphere, unless defined with an ANI threshold of 99.9997% or greater. Hence, even marine bacterial strains (if defined with a 99.5% ANI threshold as in (Rodriguez-R *et al*., 2022)), are likely to be globally distributed. Lake prokaryotes showed an intermediate level of endemism with predicted probability to be found on opposite hemispheres approaching zero for clades >99.6% ANI, which implies that analyses of lake bacteria on species level would exhibit little geographic signal, while intra-species divergence may still be affected by dispersal limitation. This would open the possibility for local adaptation, and ultimately enable allopatric speciation. The extent of dispersal limitation is thus decisive for evolution and a main theme in the present study.

It should be noted that genome-wide data is necessary to resolve the relevant time scales to study allopatric divergence. Molecular clocks suggest that 99.6% ANI corresponds to 10,000 – 600,000 years of divergence (Ochman and Wilson, 1987; Denef and Banfield, 2012) while the 16S rRNA gene by comparison is expected to accumulate less than one substitution per million years (Kuo and Ochman, 2009). The latter is thus of limited value for resolving allopatric divergence of freshwater bacteria. However, conspecific bacterial genomes from distant lakes are scarce (Louca, 2021), and therefore also biogeographic insights. To overcome this data paucity, we leveraged publicly available datasets and newly sequenced freshwater metagenomes from around the world. Read mapping against a selected set of reference genomes allowed us to identify globally abundant freshwater bacterial species and analyze their population structure. To further place the observed scales of geographic divergence in context, we compared these observations to temporal divergence within individual lakes and finally propose a conceptual model to explain contrasting patterns of intra-population diversity and dynamics.

### Definition of terms that are frequently used with different meanings elsewhere

Allele: nucleotide variant at a given locus

Habitat: Lake, pond, or other freshwater sampling site

Locus: a single nucleotide position in the genome

Polymorphism: a locus with two or more alleles

Population: conspecific organisms co-occurring in a habitat

Species: group of organisms sharing >95% ANI (genomospecies)

### Species abbreviations

F. sp.: *Ca*. Fonsibacter sp. (reference genome is a SAG), affiliated with the LD12 group

F. ubi.: *Ca*. Fonsibacter ubiquis, affiliated with the LD12 group

M. uni.: *Ca*. Methylopumilus universalis

N. abu.: *Ca*. Nanopelagicus abundans, affiliated with the acI group

*P. fin*.: *Polynucleobacter finlandensis*

*P. pan*.: *Polynucleobacter paneuropaeus*

P. ver.: *Ca*. Planktophila vernalis, affiliated with the acI group

## Results and discussion

### Detection of abundant and globally distributed freshwater bacterial species

A set of 1952 lentic freshwater metagenomes was used to screen for the 20 initially selected reference genomes, representing 20 different genomospecies (<95% ANI for all pairs) associated with the five genera *Ca*. Fonsibacter (*Alphaproteobacteria*, LD12 group), *Ca*. Methylopumilus (*Betaproteobacteria*), *Ca*. Nanopelagicus (*Actinomycetota*, acI group), *Ca*. Planktophila (*Actinomycetota*, acI group) and *Polynucleobacter* (*Betaproteobacteria*). Seven of the species were detected with high enough coverage (see Methods for used cutoffs) at a broad enough geographic range to be included in further analyses (Table 1, Fig. 1, Suppl Fig. S1), leaving a combined dataset of 225 metagenomes (Suppl. Table S1) with 10 – 111 metagenomes per species (Table 1).

**Table 1.** Species and underlying data analyzed in this study.

| Species | Ref.* | Abbr. | Reference genome/strain | Reference genome source habitat | Reference genome NCBI accession | Reference genome size (bp) | # Analyzed metag. | # Analyzed habitats | # Analyzed loci | # Polymorphic loci |
| --- | --- | --- | --- | --- | --- | --- | --- | --- | --- | --- |
| *Ca. Fonsibacter* sp. | 1 | F. sp. | AAA028-D10 (SAG) | Lake Mendota | GCA_000510845.1 | 925141 | 23 | 7 | 296334 | 7459 |
| *Ca. Fonsibacter ubiquis* | 1, 2 | F. ubi. | LSUCC0530 | Lake Borgne | GCF_002688585.1 | 1160202 | 19 | 11 | 384503 | 6242 |
| *Ca. Methylopumilus universalis* | 3 | M. uni. | MMS-RIV-30 | Rimov reservoir | GCF_006364215.1 | 1268083 | 38 | 9 | 539371 | 10365 |
| *Ca. Nanopelagicus abundans* | 4 | N. abu. | MMS-IIB-91 | Lake Zurich | GCF_002288305.1 | 1161863 | 10 | 3 | 508148 | 11124 |
| *Ca. Planktophila vernalis* | 4 | P. ver. | MMS-IIA-15 | Lake Zurich | GCF_002288185.1 | 1364004 | 13 | 5 | 430975 | 13209 |
| *Polynucleobacter finlandensis* | 5 | *P. fin.* | MWH-Mekk-B1 | Lake Mekkojärvi | GCF_018881755.1 | 2280072 | 111 | 14 | 905581 | 19467 |
| *Polynucleobacter paneuropaeus* | 6 | *P. pan.* | UB-Kaiv-W7 | Pond Kaivoslampi | GCF_003261255.1 | 1830921 | 37 | 14 | 1106814 | 15830 |

\*References: 1, Zaremba-Niedzwiedzka et al. 2013; 2, Henson et al. 2018; 3, Salcher et al. 2019; 4, Neuenschwander et al. 2018; 5, Hahn et al. 2022; 6, Hoetzinger et al. 2019

**Fig. 1.**
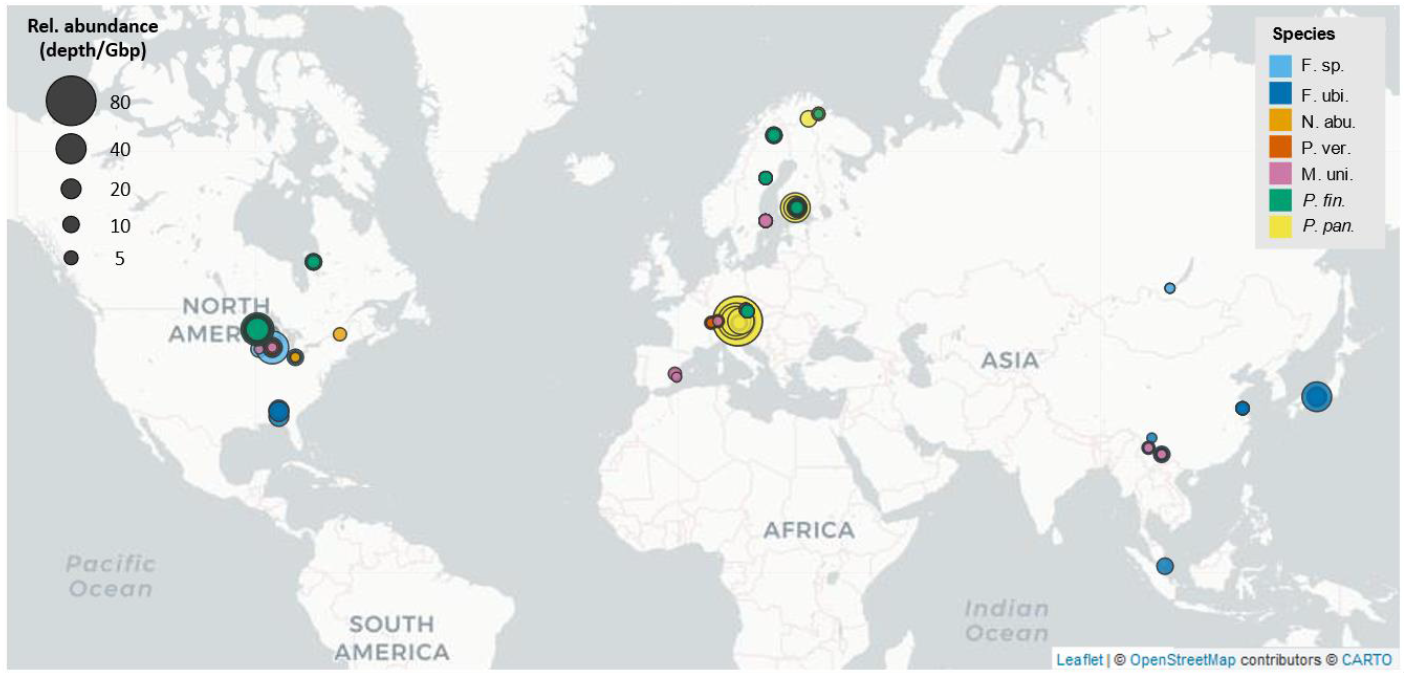
Detection of the different species (color-coded) in the used metagenomes. Each dot refers to one metagenome. Relative abundance (dot size) is given as median coverage depth of the reference genome per Gbp of metagenome (depth/Gbp) as obtained from read mapping. Note that some dots are overlaid by others (multiple metagenomes from the same site or multiple species in the same metagenome). Refer to Suppl. Fig. S1 for separate maps for each species.

Relative abundances assessed as percentage of metagenome bases mapped to a reference genome (note that this measure differs from relative cellular abundance as it depends on genome size, genome copy numbers per cell etc.) were highest for *P. pan*. in several humic ponds, with up to 13.2% mapped bases in an Alpine pond (Fig. 2). Similarly, *P. fin*. recruited up to 11.6% of bases from metagenomes of the humic Trout Bog Lake. The two species of the LD12 clade were detected at a maximum relative abundance of 6.1% in Lake Biwa (F. ubi.) and 5.4% in Lake Michigan (F. sp.). The two acI species recruited up to 1.7% in Lake Michigan (N. abu.) and 0.9% in Lake Loclat (P. ver.), and M. uni. accounted for up to 0.7% in the Rímov reservoir. Note, however, that sampling procedures (pre-filtration etc.) and sequencing methods varied for the different metagenomes. Accordingly, relative abundances are not directly comparable among metagenomes. It was still obvious that the two *Polynucleobacter* species thrived in acidic waters rich in humic matter, and they were also found to be abundant and coexisting in a few small habitats in Fennoscandia (Suppl. Table S1). In contrast, the other five species thrived in mostly larger lakes with neutral to slightly alkaline pH, where the same type of “abundant coexistence” was apparent in several lakes, with up to four of the species coexisting in Lake Zurich. While *P. pan*. and P. ver. were not detected outside Europe, at least not with high enough coverage to enable population structure to be described, the five other species covered multiple continents. F. ubi. spanned the largest geographic range of 16 200 km, from an urban water sample in Singapore to Lake Eufaula in Georgia, USA.

**Fig. 2.**
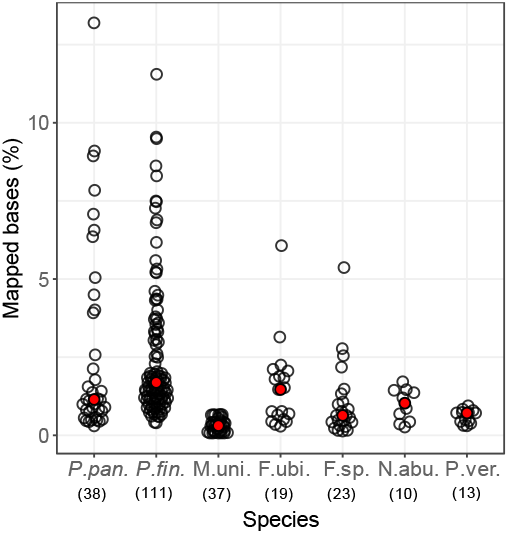
Percentage of metagenome bases mapped to the reference genomes. Each dot refers to one metagenome. Median values are shown in red. The number of metagenomes for each species is given in brackets under the x-axis.

### Species demarcation

As we define species based on DNA sequence identity in this study, we first need to discuss the feasibility and relevance of such an approach. It has been shown that bacterial genomes of isolates as well as metagenome-assembled genomes (MAGs) have a striking sparsity of ANIs between 83 and 96%. Accordingly, this gap has been used to operationally define species based on 95% ANI (Konstantinidis *et al*., 2006; Jain *et al*., 2018). The validity of this threshold has been corroborated also for natural communities when mapping metagenome reads to reference genomes (Caro-Quintero and Konstantinidis, 2012; Bendall *et al*., 2016; Garcia *et al*., 2018), ruling out potential isolation or assembly biases (Gevers *et al*., 2005). Such discernable species separated by genetic discontinuity observed in metagenomic reads were referred to as sequence-discrete populations (Caro-Quintero and Konstantinidis, 2012). In the present dataset, we observed sequence-discrete species over global scales (Fig. 3, Suppl. Fig. S2), providing ultimate support for a discontinuous diversity structure of bacteria in nature. However, the extent of diversity within species differed considerably. The two *Polynucleobacter* species appeared to be the most cohesive indicated by high ANIr_95_ (average nucleotide identity of the metagenome reads mapped to the reference genome with ≥95% identity) of 99.1% (*P. pan*.) and 98.5% (*P. fin*.) across all metagenomes used with these species (Fig. 3), accompanied by a low proportion of reads mapping within the putative “species gap” at around 90% identity in most of the metagenomes (Suppl. Fig. S2). If a peak at <95% identity appeared in the identity histogram of a metagenome, representing co-occurring *Polynucleobacter* species, these typically occurred at 80% nucleotide identity or below. Among the 111 metagenomes mapped against *P. fin*. only two from Lake Ki1 showed a minor peak at 91% identity, thus falling within the “species gap” and possibly representing a closely related sister species. No such peak occurred among the 38 metagenomes mapped against *P. pan*. These observations align well with previous observations of genome comparisons of isolated *Polynucleobacter* strains that show a dominance of strain**-**pairs with >97% and <84% ANI and only a few exceptions in between (Hoetzinger and Hahn, 2017; Hahn *et al*., 2022). The other betaproteobacterium M. uni. also appeared as cohesive species with an ANIr_95_ of 98.5%, yet, peaks of mapped reads possibly indicating co-occurring closely related species appearing at around 86% identity in several metagenomes (Suppl. Fig. S2). In contrast to the cohesive betaproteobacterial species, the two acI species were the most diverse according to ANIr_95_ values of 97.7% (N. abu.) and 97.6% (P. ver.) across all metagenomes used with these species (Fig. 3). It was evident that all blast histograms for the two acI species displayed a prominent second peak slightly below 80% identity (Suppl. Fig. S2), suggesting that related species typically co-occur together in the same habitat. A rather different pattern emerged for the two LD12 species, with a strikingly low proportion of mapped reads with <80% nucleotide identity in the metagenomes. Similarly to acI, these populations were rather diverse, indicated by broad peaks in the blast histograms. In several histograms, a second peak was apparent between 85 and 95% identity (Suppl. Fig. S2). This suggests that either multiple closely related sister species frequently co-occur, or that species boundaries are blurred for LD12. In line with that, previous comparisons of single amplified genomes (SAGs) from different lakes showed ANIs of >90% in LD12 but around 80% in acI for most inter-species comparisons (Garcia *et al*., 2018). Overall, the identity distributions of mapped metagenome reads seem to corroborate the usefulness of the commonly used 95% ANI threshold for bacterial species delineation, given the peaks in the histograms at >95% identity in all metagenomes (except for eight excluded ones, Suppl. Fig. S3) and a “species gap” somewhere between 85 and 95% for most of them. It is nevertheless important to note that intra-species divergence as well as the frequency of putative sister species occurring within the “species gap” varied considerably among taxa. Hence, the 95% ANI threshold is not equally suited for delineating all different taxa. It should rather be seen as a pragmatic (lower) limit for delineation that works surprisingly well for many taxa, but not as a token for a universal composition of intra- and inter-species divergence.

**Fig. 3.**
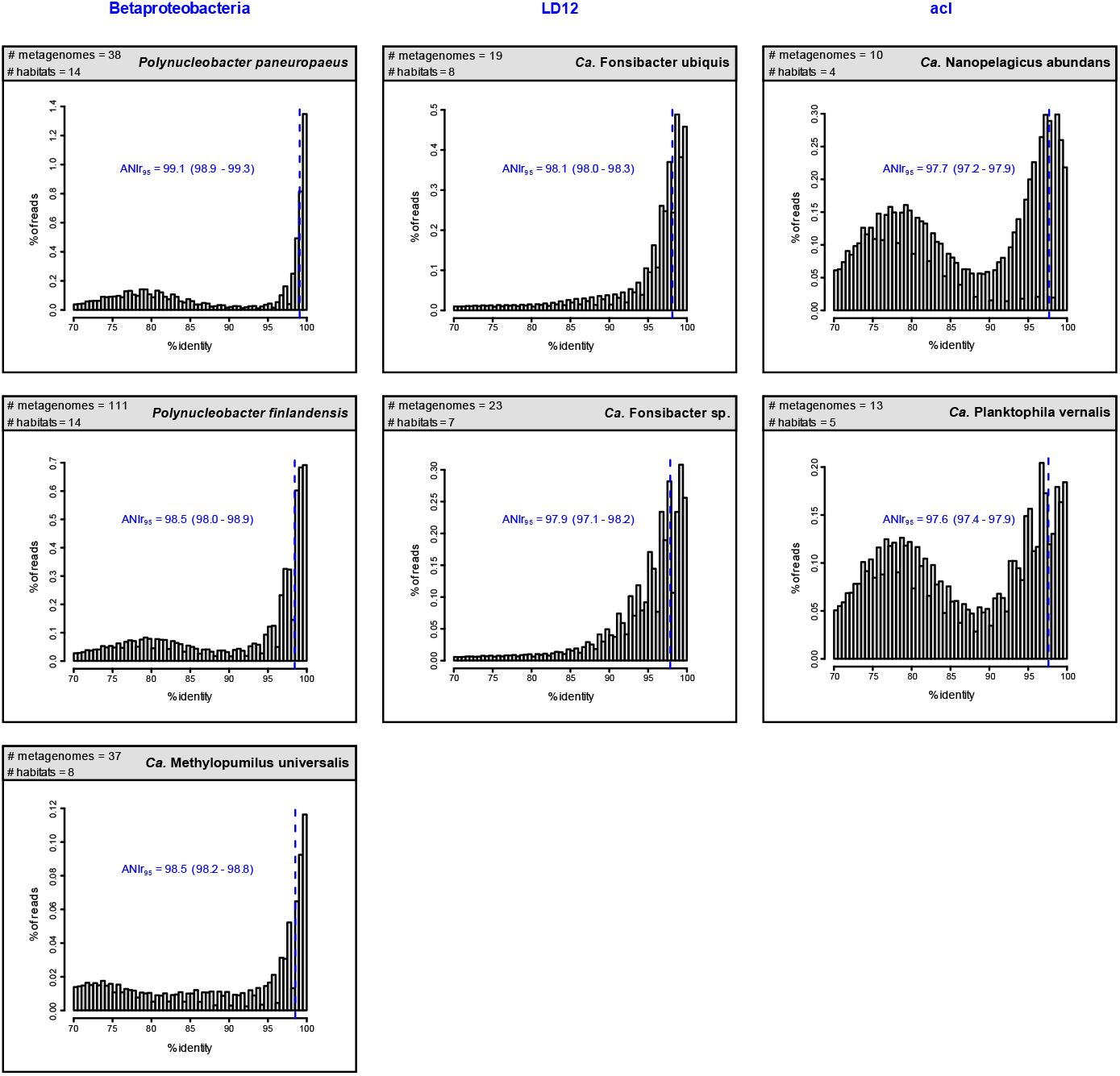
Histograms of metagenome reads mapped to the reference genomes by Blast. The numbers of metagenomes and different habitats underlying each plot are given on the upper left. For each species the ANIr_95_ value across all respective metagenomes is given and indicated by a blue dashed line. The range of ANIr_95_ values when computed for the respective metagenomes separately is given in brackets.

### Intra-species diversity

To characterize intra-species divergence beyond ANIr_95_ (Fig. 4A), we calculated other measures of genomic variation in metagenomes based on mappings with ≥95% identity thresholds (Suppl. Table S2). Interestingly, a simple measure giving the proportion of polymorphic loci (Fig. 4B) showed very similar results to the nucleotide diversity (π), giving the average number of nucleotide differences per locus (Fig. 4C). The proportion of polymorphic loci ranged from 0.47% in *P. pan*. to 2.04% in P. ver., and π from 0.12% to 0.75%. The obtained ranking of species by these metrics was nevertheless slightly different than that obtained based on ANIr_95_. Especially *P. fin*. was relatively diverse according to π but not according to ANIr_95._ A potential bias of ANIr_95_ can stem from the choice of reference genome, while π is not influenced by that choice. Theoretically, the loci considered for calculating π could change depending on the reference genome used, but as we only included loci that were sufficiently covered in all metagenomes, the analyses mainly refer to “core loci” expected to be present in almost all genomes of the species, while the more flexible part of the genomes would have been excluded. The median ANIr_95_ of *P. fin*. was indeed lower when excluding the metagenomes from Lake Mekkojärvi where the reference strain was isolated from (Suppl. Fig. S4), i.e. ANIr_95_ of *P. fin*. was elevated because many metagenomes for this species (36 out of 111) were from the home habitat of the used reference strain. However, even when excluding these metagenomes, *P. fin*. still shows the second lowest diversity according to ANIr_95_ (Suppl. Fig. S4). Its disproportionally high π suggests a distinctive diversity structure compared to the other species. Besides the discussed diversity measures it would thus be interesting to know the strain diversity, especially for analyzing population dynamics (see below). This would help to clarify if abundance dynamics of different strains are uncoupled, e.g. due to environment-dependent fitness differences that may also include strain-specific differences in predation susceptibility. Strain diversity is expected to correlate with nucleotide diversity, yet, a given nucleotide diversity could be realized by a higher number of closely related strains or by a lower number of distantly related strains. To calculate a diversity metric that represents strain diversity better than π, we used Shannon entropies of polymorphic loci with non-correlated allele frequencies (H_noncorr_). We benchmarked our method to compute H_noncorr_ as a proxy for strain diversity on a simplified dataset, which showed that it reflected strain diversity better than π (Suppl. Fig. S5, Suppl. Table S3). In the natural data it roughly divided the seven species into four groups of increasing diversity (*P. pan*., *P. fin*. + M. uni., F. ubi. + F. sp. and N. abu. + P. ver.), whereby *P. pan*. showed by far the lowest median H_noncorr_ (Fig. 4D).

**Fig. 4.**
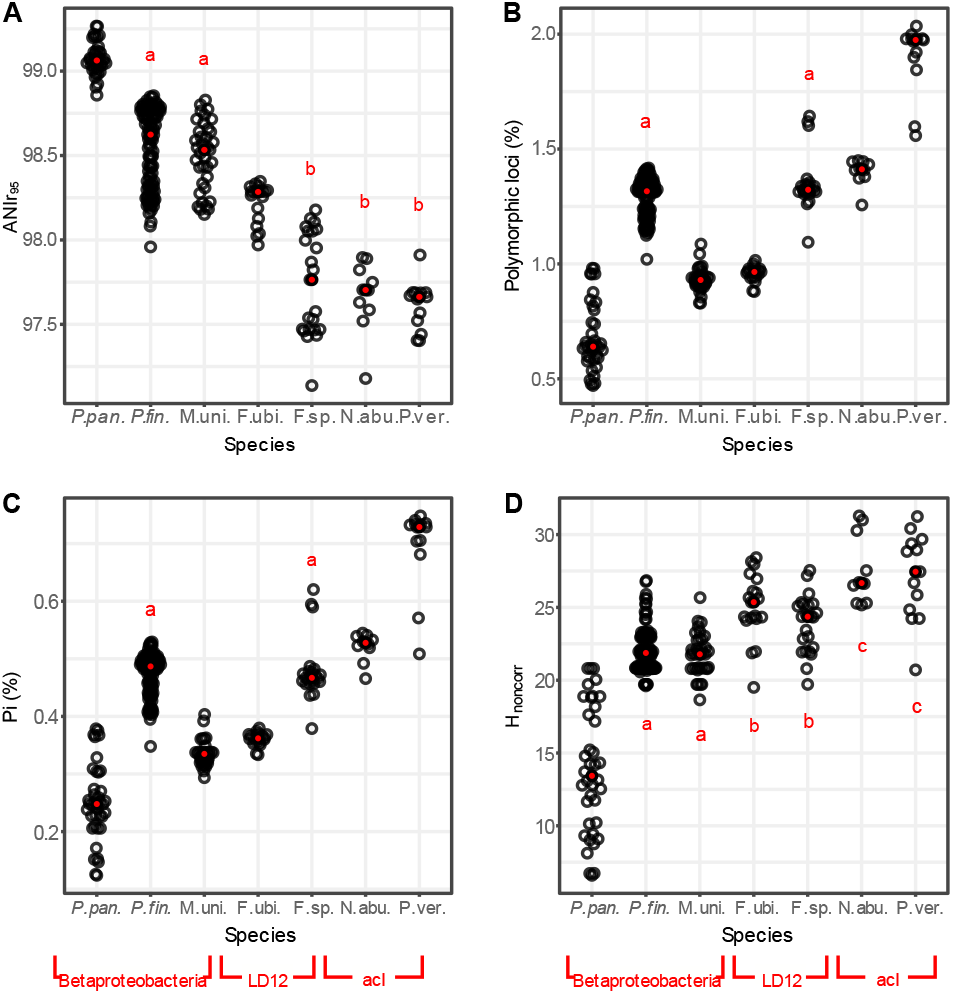
Intra-population diversities for the seven species according to different metrics. Each dot refers to one metagenome. Median values are shown as red dots. Pairs that are not significantly different (p >0.05) according to Wilcoxon-Mann-Whitney Rank Sum test are marked with a common letter.

Note that this study covers allelic diversities within the core genome, although intra-species diversity in prokaryotes can to a large extent stem from gene content diversity, i.e. the accessory genome. The implications of the accessory genome on the evolution of prokaryotic species are certainly substantial, yet, beyond the scope and reach of this study.

### Homologous recombination

Linkages between nearby alleles in the genome give hints about homologous recombination rates, although other factors such as selection may influence these linkages too. Loci are in linkage disequilibrium (Lewontin and Kojima, 1960) when the association of their allele frequencies are non-random. Nearby loci are more likely to be transferred together in recombination events and are thus expected to show linkage disequilibrium in recombinogenic bacteria. Linkage measured as the normalized squared correlation coefficient r^2^_norm_ was highest in *P. pan*., up to 0.7 between neighboring alleles (Fig. 5). A maximum value of 1 would mean that pairs of alleles at polymorphic loci with a given distance always occur together on the same read or read pair. Linkage approached a baseline around 350 bp, suggesting that the sequence length of most homologous recombination events was shorter than that (cf. (Ansari and Didelot, 2014)). The other six species showed lower linkage disequilibrium, with maximum r^2^_norm_ values ranging from 0.3 (F. sp.) to 0.5 (*P. fin*.) and r^2^_norm_ versus distance plateauing earlier, between 100 bp (N. abu.) and 300 bp (*P. fin*.). Species coherence in *P. pan*. was suggested earlier to be promoted by high homologous recombination rates (Hoetzinger *et al*., 2021). A clear relationship between species coherence (Fig. 3) and putative recombination indicated by linkage (Fig. 5) was however not apparent for the other species. Species coherence may also stem from other causes such as periodic selection (Atwood *et al*., 1951) or the K/θ ≥ 4 rule derived from population genetic theory (Birky and Maughan, 2021). Although the discontinuity below 95% ANI seems to be a rather universal feature in bacteria, coherence in different species may be shaped by different microevolutionary processes and homologous recombination may not be obligatory for species clustering.

**Fig. 5.**
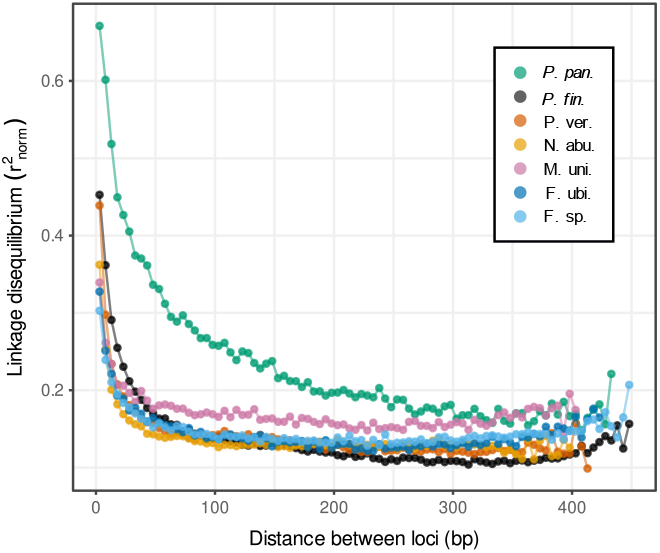
Linkage between polymorphic loci. The normalized correlation coefficient between pairs of loci (r^2^_norm_) was averaged over different pairs with similar distance between the loci, using a 5 bp window. Only windows with at least 20 values were plotted. Higher r^2^_norm_ values suggest that loci at the given distance were more likely inherited together.

### Divergence with spatial distance

To assess the impact of geographic separation on population differentiation, we analyzed fixation indices (F_ST_) from pairwise comparisons of populations versus spatial distance between the respective sampling sites (Fig. 6). F_ST_ is a measure of population divergence that relates inter- to intra-population diversity. For instance, an F_ST_ of 0 would mean that the diversity when comparing sequences from different populations would be the same as the diversity within sequences of the same population. The maximum F_ST_ of 1 would mean that all polymorphic loci have a fixed allele in one population and an alternative allele in the other population. All species showed a significant increase in F_ST_ with spatial distance according to Mantel tests. Interestingly, the increase appeared rather linear with distance for most of them. This may at first seem surprising if one considers that dispersal when modeled as a diffusion-like process decreases with the square of the distance (Louca, 2021). Possibly, long-range dispersal could be governed by incremental dispersal events between nearby sites rather than direct dispersal between distant sites (stepping-stone dispersal), which would explain the more linear relationship observed. Yet, the slope tends to be higher at shorter distances (Suppl. Fig. S6), which may reflect a more quadratic decay where direct dispersal between sites is more relevant. We have previously observed a similar pattern when analyzing genome similarities of *P. pan*. isolates (Hoetzinger *et al*., 2021). The observation from that earlier study, that divergence of *P. pan*. does not increase with spatial distance when only longer distances (> few hundred km) are considered, was also corroborated in the present work. When comparing the rate of F_ST_ increase with spatial distance, it is conspicuous that six of the species were in a similar range (0.039 – 0.114 ΔF_ST_/1000km), while F. ubi. showed a substantially lower divergence (0.0024 ΔF_ST_/1000km). Interestingly, the type strain of this species was isolated from the brackish coastal lagoon Lake Borgne (salinity of 0.24%) and could be cultivated at salinities up to 0.47% (Henson *et al*., 2018). It is conceivable that higher stress tolerance or a broader niche range of F. ubi. allows the species to better survive dispersal and accordingly explains its low geographic divergence. We do not currently know if it is able to survive a stopover in the oceans to disperse more effectively between continents, but even for the other species, the North Atlantic and North Pacific Ocean did not seem to pose strong dispersal barriers as no offset to higher F_ST_ values was observed for trans-oceanic relative to intra-continental comparisons (Fig. 6). Between Asia and North America, the Bering Strait might facilitate dispersal. Yet, the population structures of M. uni. and F. sp., with metagenomes analyzed from Asia, North America, and Europe, did not show any disproportionally high population divergence between North America and Europe, which are not well connected through inland waters (Fig. 7). It should be noted though that many factors potentially influence microbial dispersal across oceans. It has been shown recently that terrestrial and dust-associated bacteria were more prevalent in the atmospheric community over the Atlantic compared to over the Pacific Ocean (Lang-Yona *et al*., 2022), which might hint at facilitated dispersal of continental bacteria across the Atlantic. Dust particles are known vectors for bacterial dispersal, and e.g. desert dust clouds from Africa can be transported to North America (Griffin 2007). Freshwater bacteria may also hitchhike with migratory waterfowl that cross oceans (Watts *et al*., 2021; Conklin *et al*., 2022; Piersma *et al*., 2022). It remains unclear which modes of trans-oceanic dispersal are most relevant. Anyhow, our results suggest that the North Atlantic and North Pacific Ocean are not particularly strong dispersal barriers for the abundant freshwater bacteria analyzed in this study.

**Fig. 6.**
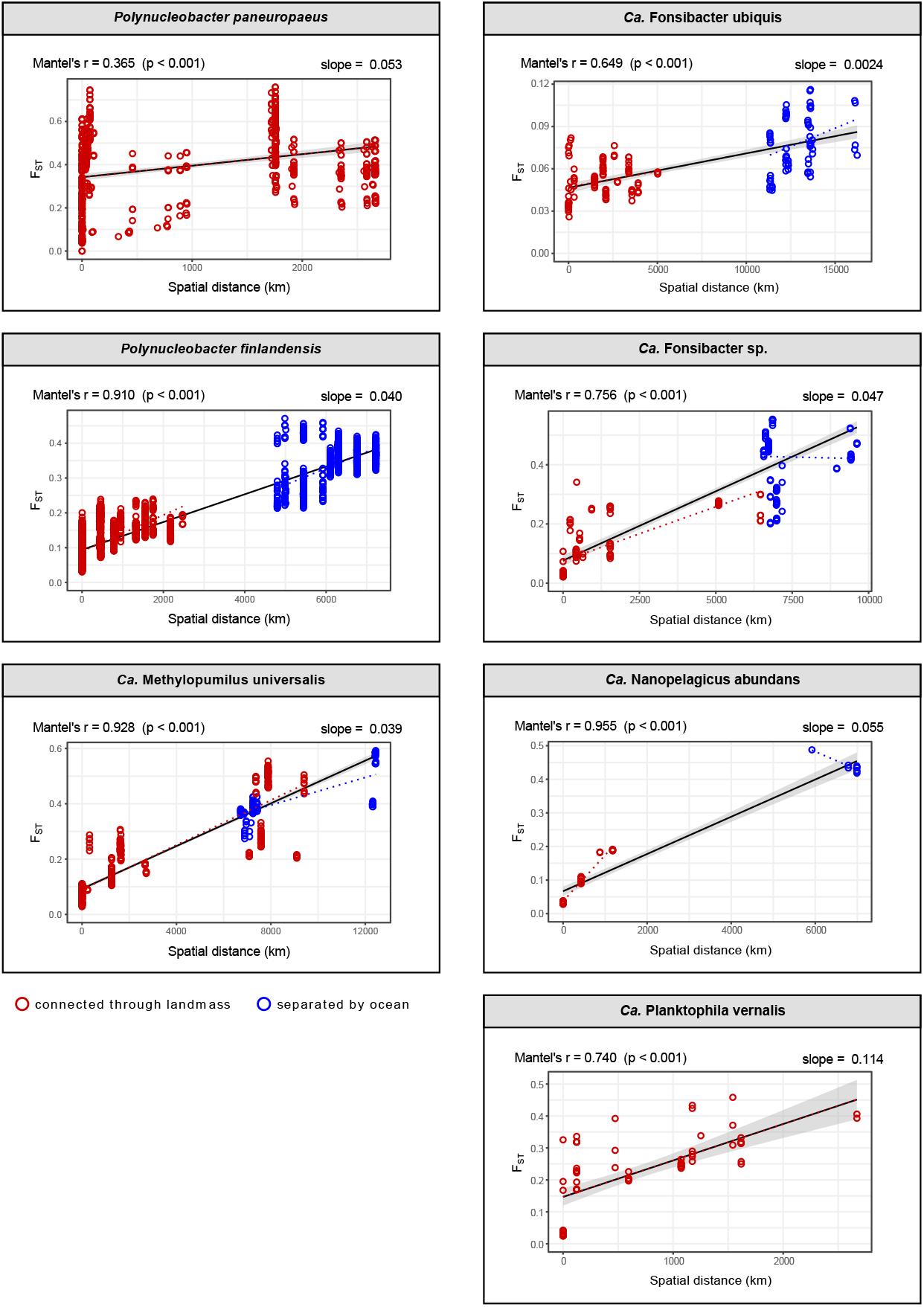
Population differentiation (F_ST_) versus spatial distance between habitats for the seven investigated species. Red dots refer to comparison within Eurasia or North America, blue dots refer to comparisons between Eurasia and North America. The latter are not markedly shifted towards higher F_ST_ values, which suggests that oceans did not pose strong dispersal barriers. Linear regressions across the whole distances are shown as black lines. The respective slopes (ΔF_ST_ / 1000 km) are given on the upper right of each plot. Linear regressions for only red and only blue datapoints are shown in the respective color as dotted lines. Spearman rank correlations across the whole distances assessed through Mantel tests are given on the upper left of each plot.

**Fig. 7.**
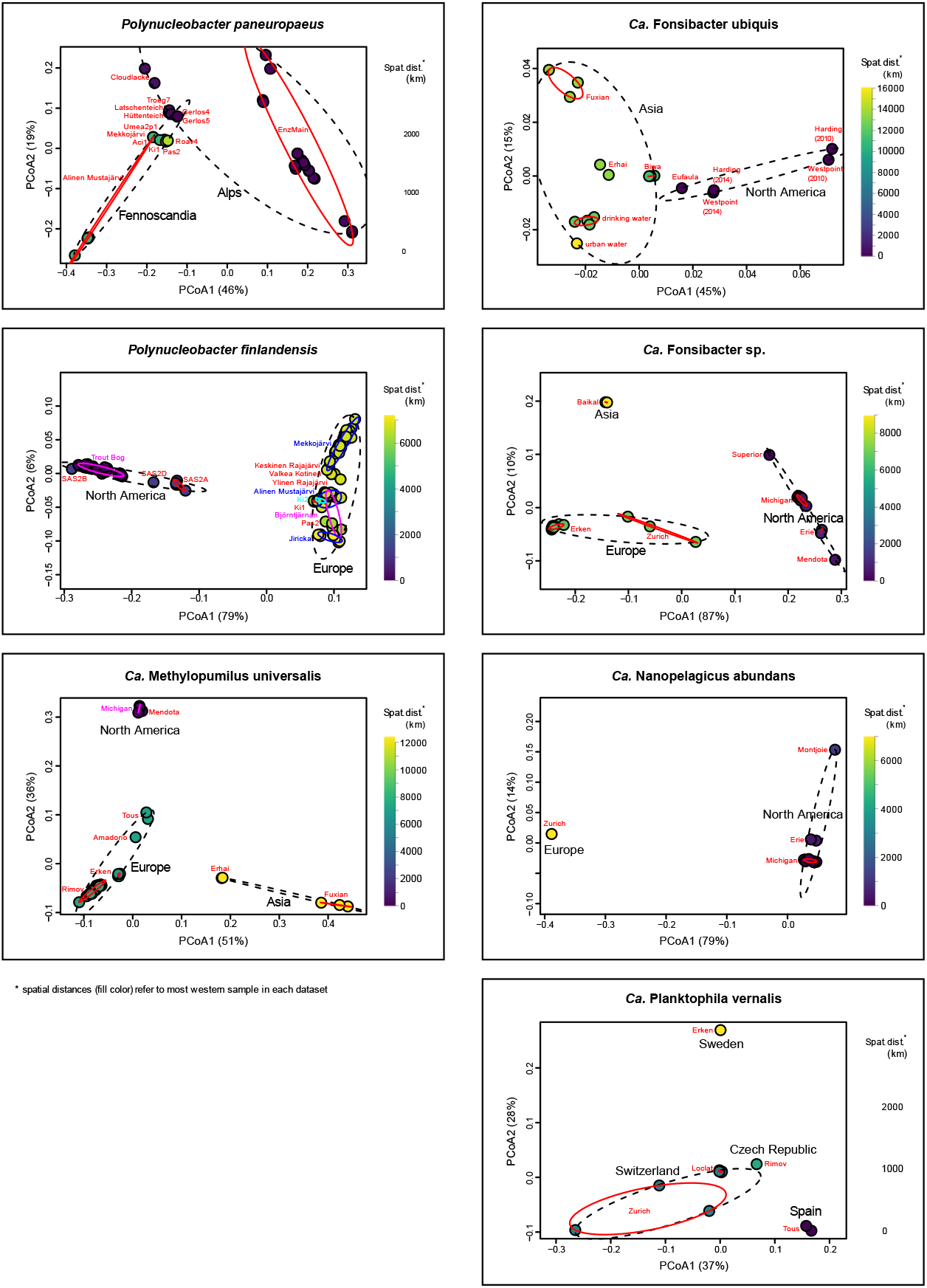
Principal coordinates analyses of F_ST_ values. Each dot represents one metagenome. Spatial distances to the most western metagenome for each species are visualized by the color gradient. Metagenomes from the same continent, region or country are edged by black dotted ellipses. Metagenomes from the same habitat are edged by red ellipses. Habitat names are given in colored font. The percentage of variation explained along each axis is given in the axis titles.

### Divergence within habitats

To put the observed population divergence with geographic distance into perspective, we analyzed spatial and temporal divergence also within habitats. First, we assessed population differentiation along the water column by analyzing metagenomes sampled on the same day but from different depths (Fig. 8). Most comparisons (92%) did not show pronounced differentiation (F_ST_ < 0.1). Strongest divergence (F_ST_ = 0.20) was found in P. ver. between the epilimnion (5m) and hypolimnion (40-80m) of Lake Zurich in November 2015. *P. fin*. showed differentiation with F_ST_ values up to 0.16 between samples from oxic surface water (0.25m) in Lake Björntjärnen as compared to anoxic bottom water (7m) from the same lake, both collected in September 2018. Overall, divergence across the water column appeared to be minor relative to geographic divergence.

**Fig. 8.**
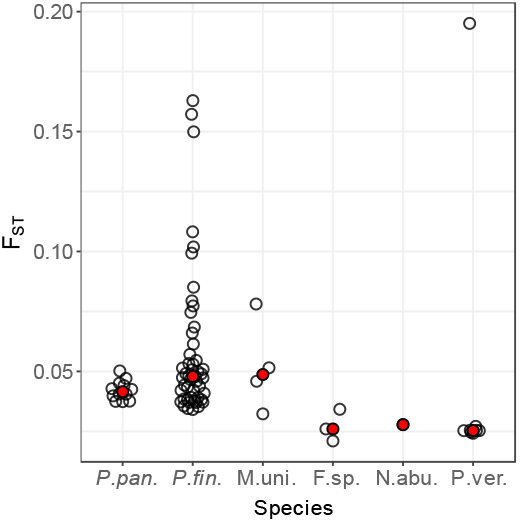
Population differentiation within the water column. F_ST_ between samples taken from the same habitat at the same time but from different depths. Median values are shown in red.

For four species in five different habitats there were extended metagenomic time series (up to 7 years) available (Bendall *et al*., 2016; Mehrshad *et al*., 2018; Andrei *et al*., 2019; Kavagutti *et al*., 2019; Mondav *et al*., 2020; Buck *et al*., 2021), enabling us to study population differentiation over time. As we had observed for spatial distance, F_ST_ increased significantly also with time (Fig. 9). Yet, the variability in the rate of this increase was much higher than for geographic divergence. The rate ranged from 0.0065 to 0.37 ΔF_ST_/1000days for the different time series, as compared to a much narrower range of 0.039 to 0.053 ΔF_ST_/1000km spatial distances for the same four species. The large variability in temporal divergence points to fundamentally different population dynamics among different species, or possibly also among different populations within the same species. Changes in relative abundance and intra-population diversity with time are shown in Suppl. Fig. S7. The two extremes in terms of temporal dynamics were observed between F. sp. in Lake Erken, which showed very slow and steady divergence with a maximum F_ST_ of 0.043 in seven years, and *P. pan*. in the Alpine pond EnzMain, with an F_ST_ up to 0.54 between two samples retrieved only 2 months apart. Note for comparison that maximum F_ST_ values between different habitats were 0.55 and 0.76 for F. sp. and *P. pan*., respectively. Allele frequency distribution changes over time provided some hints towards potential explanations for this contrasting divergence rates.

**Fig. 9.**
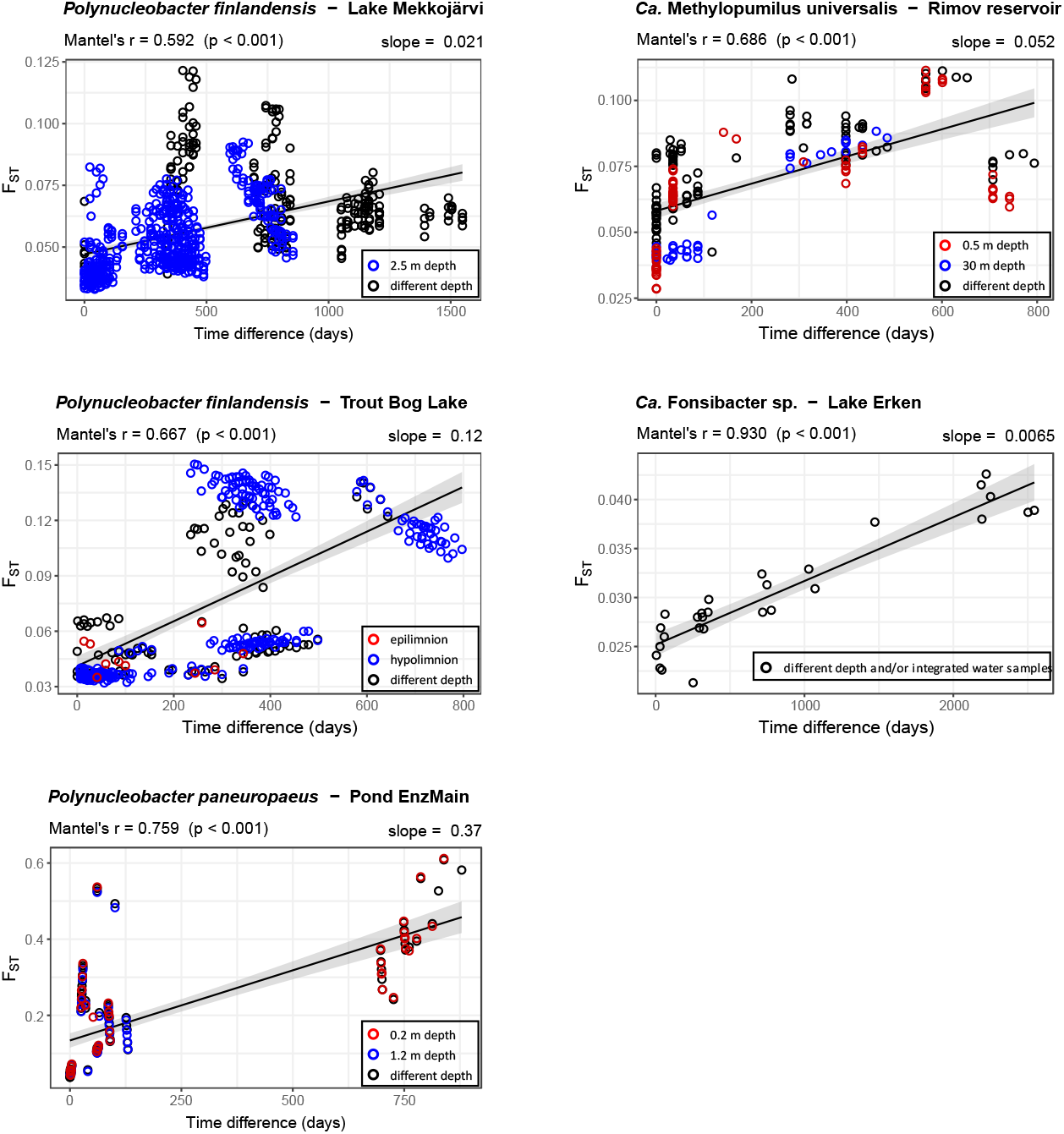
Population differentiation (F_ST_) versus time difference between sampling dates for five time series datasets. Colored dots refer to comparisons between samples taken from the same depth. These are not markedly shifted towards lower F_ST_ values, which suggests that population differentiation across the water column is minor overall. Linear regressions are shown as black lines. The respective slopes (ΔF_ST_ / 1000 days) are given on the upper right of each plot. Spearman rank correlations assessed through Mantel tests are given on the upper left of each plot.

### Intra-population dynamics

The F. sp. allele frequency distribution in Lake Erken hardly changed with time, while it was highly dynamic in *P. pan*. in Pond EnzMain (Suppl. Fig. S8). In the *P. pan*. population, periods when polymorphic loci were dominated by one allele alternated with periods when major and minor alleles were present in similar proportions. The former might indicate one dominating subpopulation, while the latter might represent two subpopulations present in similar abundance. To see which alleles show synchronous changes over time and might thus be associated with the same subpopulation, we calculated a correlation matrix for all alleles based on their allele frequencies in the different metagenomes of the time series (Fig. 10A). While alleles seemed to cluster into different groups in *P. pan*., no clear clustering was apparent in F. sp. After hierarchical clustering, *P. pan*. alleles were grouped into nine and F. sp. into eleven clusters with coherent dynamics (Suppl. Fig. S9). Note that the number of groups depends on the clustering criterium used and is thus somewhat arbitrary, and that the distinction between clusters was weak in F. sp. as mentioned above. By tracing the relative abundances of groups over time, we could thus illustrate the distinct population dynamics between *P. pan*. and F. sp. (Fig. 10B). In *P. pan*., two subpopulations appeared to alternately dominate the population in the pond. Dynamics of the third largest allele group (green) were concurrent to that of the largest (red) in 2020, but not in 2018 when the former had low relative abundance. The groups could possibly be considered as associated to the same subpopulation in 2020, and it is conceivable that the alleles of the “green group” were introduced into that subpopulation by homologous recombination or simply through mutation and eventually became fixed in the subpopulation. In that context, it is worth noting that alleles are not necessarily linked to certain subpopulations in the long run but can become unlinked through high homologous recombination rates as for *P. pan*. (see above). In any case, allele frequency dynamics in the *P. pan*. population were remarkable in that dominant alleles were swept from the population only to become dominant again at later times. For example, 117 out of 3977 alleles that were absent (allele frequency = 0/15) in May 2018, became the major allele (frequency ≥ 8/15) in June 2020, were then absent in July 2020, and became the major allele again in September 2020. Sweeps were thus incomplete and transitory in *P. pan*., i.e. most alleles probably remained persistently in the population, albeit at times below the detection limit. It seems less likely that alleles were completely lost and reestablished repeatedly through either novel mutation or recolonization. These observations are in line with the constant-diversity model (Rodriguez-Valera *et al*., 2009), where a complete sweep of diversity by an over-dominant strain is prevented by strain-specific phage predation constraining and counteracting “blooms” of individual strains (i.e. “kill the winner” (Thingstad, 2000)). It remains to be tested whether or not phage predation was responsible for the observed dynamics in *P. pan*., and it cannot be excluded that complete selective sweeps leading to periodic selection (Guttman and Dykhuizen, 1994) are happening over longer timescales. However, high homologous recombination rates in *P. pan*. are supposed to unlink different genes in the genome, and putative sweeps might thus be gene-specific rather than genome-wide (Shapiro and Polz, 2014). Sweeps of diversity and the homogenizing effect of homologous recombination could both constrain diversification and potentially explain the overall low nucleotide diversity in *P. pan*. In contrast to *P. pan*., the F. sp. population showed an exceptionally slow and continuous evolution with time. Nucleotide diversity varied only marginally from 0.46 to 0.47% throughout the time series (Suppl. Fig. S7) and there were no apparent subpopulations with distinct dynamics (Fig. 10, right). The small variation in the F. sp. compared to the *P. pan*. population is remarkable considering the longer timespan of the former dataset (7.0 vs. 2.4 years). The F. sp. population could be viewed either as a consortium of a large number of strains that stably coexist, or equally well as one diverse but coherent population. A factor that may have importantly contributed to the striking differences in intra-population diversity and dynamics is population size. The surface area of Lake Erken is about 14,000 times larger than that of Pond EnzMain, so we can conclude that the F. sp. population in the former is much larger than the *P. pan*. population in the latter. A larger population is expected to sustain a larger diversity and be less susceptible to changes in diversity.

**Fig. 10.**
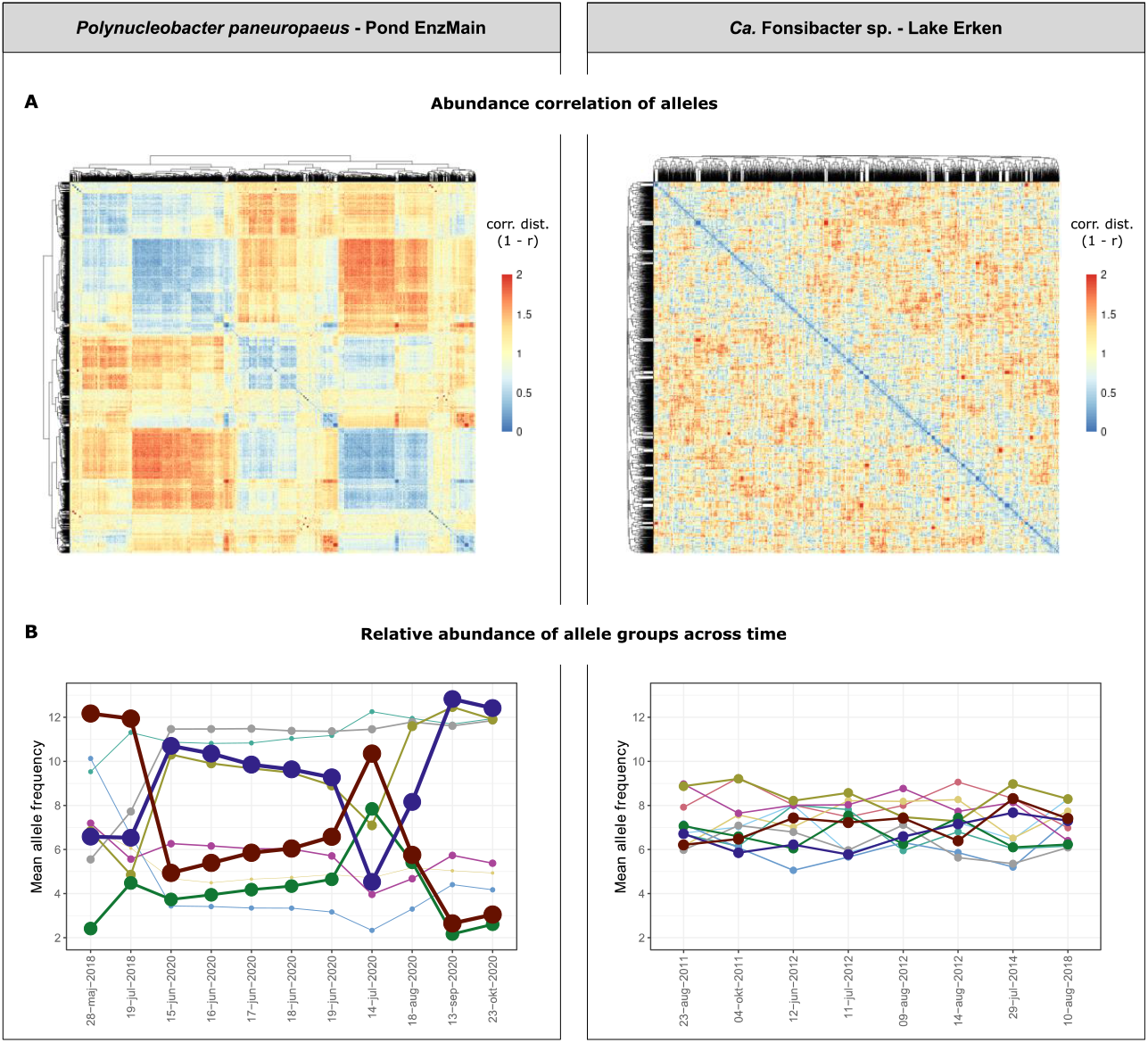
Allele frequency dynamics for two example species with contrasting patterns. **(A)** Clustering of alleles based on spearman rank correlation coefficients of allele frequencies across time. **(B)** Alleles were clustered into nine (*P. pan*.) and eleven (F. sp.) groups based on the correlations shown in A (see also Suppl. Fig. S9). The plots show the abundance of each group of alleles (mean allele frequency) across time. Dot size, line size and color density are proportional to the number of alleles in the group. Note that the timepoints are not equidistant and that the Lake Erken series spans a significantly longer time than the Pond EnzMain series.

Overall, we can formulate two contrasting conceptual models of microevolution (Fig. 11). On the one hand, recurring clonal expansions, with different genotypes alternately dominating the population. On the other hand, slow continuous microevolution, where a high diversity of genotypes persists at constant relative abundances. The type of dynamics followed by different populations of a species may have crucial implications for its evolution, e.g. its potential for diversification or maintaining diversity. Further studies including multiple populations of the same species in different habitats and with different population sizes might clarify if the type of dynamics tends to be species-specific, or rather depends on habitat-related factors such as population size or the set of genotypes present in a particular population. Such insights may help us understand the processes governing evolution in different bacterial taxa and eventually clarify how the vast prokaryotic diversity that has been uncovered through DNA sequencing is maintained by nature.

**Fig. 11.**
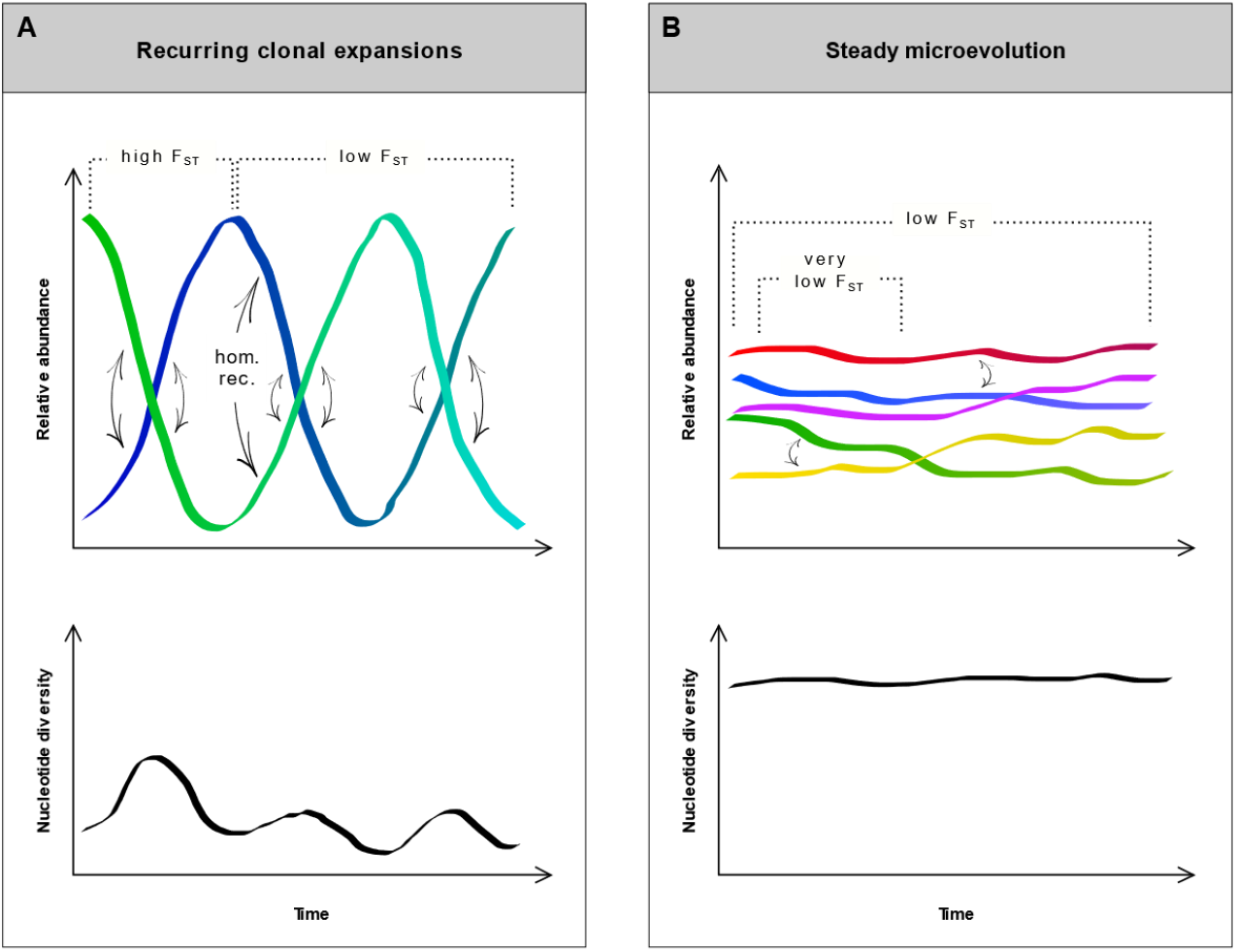
Two conceptual models based on two putative modes of microevolution observed in this study. **(A)** Strains quickly rise and fall in abundance, and the population is mostly dominated by only one strain at any given timepoint. Differentiation (F_ST_) between populations close in time (as well as between populations of nearby habitats) can thus be exceptionally high. Nucleotide diversity within the population varies strongly. It is particularly low when one strain is dominating and higher during times when multiple strains are present at similar abundances. However, due to the low number of strains present in the population nucleotide diversity is generally low and may even be reduced through extensive homologous recombination (black arrows), which counteracts divergence between strains. **(B)** A high diversity of strains stably coexists in the population. Differentiation (F_ST_) steadily but slowly increases with time. Nucleotide diversity is maintained at a constantly high level.

## Conclusion

All investigated species formed sequence-discrete populations across the studied geographic scales, with peaks of mapped reads at >95% identity, corroborating the usefulness of the widely used 95% ANI threshold for species delineation. Still, divergence within species differed considerably, pointing at distinct microevolutionary mechanisms that shape the diversity within different taxa. Population differentiation increased with spatial distance in all species, although major dispersal barriers were not apparent and oceans did not seem to pose strong dispersal barriers. Species with broad habitat ranges may be dispersed particularly well, as suggested by minimal geographic divergence of the salt-tolerant *Ca*. Fonsibacter ubiquis. Population structuring along water column depth gradients was mostly minor. In contrast, we observed striking differences between populations in their temporal dynamics. While *Polynucleobacter paneuropaeus* from an Alpine pond reached a similar divergence as seen over pan-European scales within only two months, the divergence of a *Ca*. Fonsibacter sp. population in Lake Erken was considerably lower over a seven-year period than that seen between any two populations from different lakes. We thus propose two contrasting models to explain microevolution (I) recurring clonal expansions resulting in low-diversity populations at any given time point as observed in *Polynucleobacter paneuropaeus* versus stable progression of diverse populations as seen in the Fonsibacter population.

## Materials and methods

### Reference genomes

The 20 reference genomes, 18 from isolated strains and 2 SAGs, were characterized by ANI <95% for all pairwise comparisons (Suppl. Table S4). They were chosen based on several publications that detected the respective species in different habitats with high relative abundance or across a broad geographic range (Henson *et al*., 2018; Neuenschwander *et al*., 2018; Salcher *et al*., 2019; Hahn *et al*., 2021, 2022; Hoetzinger *et al*., 2021). They were associated to five different genera (*Ca*. Fonsibacter, *Ca*. Methylopumilus, *Ca*. Nanopelagicus, *Ca*. Planktophila and *Polynucleobacter)*. The selection of reference genomes is biased by personal preference of the first author *(e*.*g. Polynucleobacter* genomes are overrepresented). The seven finally analyzed species should thus not be regarded as a fair representation of the most abundant or prevalent bacteria in global freshwater ecosystems but as a rather arbitrary yet widespread and abundant subset of those.

### Metagenomes

NCBI SRA was screened for aquatic metagenomes by querying Illumina whole genomes sequenced datasets with any of the keywords: “water metagenome”, “peat metagenome”, “seawater metagenome”, “marine metagenome”, “marine plankton metagenome”, “marine sediment metagenome”, “karst metagenome”, “lagoon metagenome”, “lake water metagenome”, “glacier lake metagenome”, “freshwater metagenome”, “freshwater sediment metagenome”, “aquatic metagenome”, “bog metagenome”, “drinking water metagenome”, “estuary metagenome”, “Winogradsky column metagenome”, “alkali sediment metagenome”, “sediment metagenome” or “ground water metagenome”. We used a custom script (*filtering_and_converting_coordinates*.*py)* to filter this table for lentic freshwater metagenomes with associated GPS coordinates. The script used keywords to exclude (e.g. sediment, viral, river) and filtered for the string ‘fresh’ in the metagenome metadata, and further checked for entries in various columns containing geographic coordinates. The initial table of 21 806 metagenomes was thereby reduced to 1646 metagenomes. We should note that the aim was not to screen public metagenomes exhaustively and that our script likely removed metagenomes that were sampled from lentic freshwater habitats and for which GPS coordinates could have been obtained by more elaborate screening. We complemented these 1646 metagenomes with the 273 metagenomes of the stratfreshDB (Buck *et al*., 2021). Besides those previously published metagenomes, we generated 33 metagenomes (Suppl. Table S5) for this study, amounting to 1952 lentic freshwater metagenomes used to screen for the target species. The newly generated metagenomes were obtained from a *P. pan*. -specific selection of humic lakes and ponds in the Austrian Alps and Northern Lapland. The selection was based on high observed relative abundances of *P. pan*. in an amplicon dataset of a protein-encoding gene (Hahn *et al*., 2021) and detection based on isolated genomes (Hoetzinger *et al*., 2021). Fourteen metagenomes were sampled in 2018 and nineteen in 2020, and a total of twenty are part of a time series from Pond EnzMain in the Austrian Alps. The 33 samples (400 – 700 ml each) were sequentially vacuum filtered through 0.8 μm and 0.2 μm Whatman Nucleopore filters (47 mm diameter) to enrich for the relatively small *P. pan*. bacteria (Hoetzinger *et al*., 2019) on the latter filters. For both 0.8 and 0.2 μm filtration, two to four filters were used for each sample as they typically clogged after passing through ∼150 ml of the humic waters. DNA was extracted from only the 0.2 μm filters for the 2018 samples, but from both 0.8 and 0.2 μm filters for the 2020 samples. DNA was extracted using phenol-chloroform-isoamyl alcohol as previously described (Schauer 2003). The 0.8 and 0.2 μm fractions of the metagenomes from 2020 were sequenced separately and have different SRA accessions in NCBI but were combined for the analyses in this paper to obtain higher coverage, as *P. pan*. was also detected in the 0.8 μm fractions. Details about the samples, library preparation and sequencing methods are given in Suppl. Table S5.

### Pre-selection of metagenomes/species

To avoid a computationally intensive mapping of all metagenomes against all reference genomes, we first filtered the metagenomes on genus level using taxonomic classification of assembled MAGs. Here, metagenomes were assembled using MEGAHIT v1.2.9 (Li *et al*., 2015), after downloading from SRA with parallel-fastq-dump (https://github.com/rvalieris/parallel-fastq-dump) and quality filtering with fastp v0.20.1 (Chen, 2023). MAGs were clustered out with MetaBAT 2 v2.15 (Kang *et al*., 2019) after mapping reads with Bowtie 2 v2.4.4 (Langmead and Salzberg, 2012) and obtaining coverage data with SAMtools v1.10 (Danecek *et al*., 2021). Scripts used for this can be found at (https://github.com/moritzbuck/SRAnsack). Completeness and contamination of the MAGs were estimated with CheckM v1.2.0 (Parks *et al*., 2015) and MAGs more than 30% complete and less than 3% contaminated where taxonomically classified using GTDB-Tk v2.1 (Chaumeil *et al*., 2022) with the GTDB release 207 (Parks *et al*., 2022). To further pre-filter the obtained set of metagenomes on species level, a fraction of each metagenome was mapped against the reference genomes. To this end, the first 10% of reads from the fastq files were downloaded using fastq-dump from the SRA Toolkit (Leinonen *et al*., 2011) and mapped against the reference genomes using Bowtie 2 (Langmead and Salzberg, 2012) with a 95% identity cutoff (settings: --ignore-quals --mp 1,1 -- rdg 0,1 --rfg 0,1 --score-min L,0,-0.05). Average coverage depth across the reference genome was computed using SAMtools (Danecek *et al*., 2021), BEDTools (Quinlan and Hall, 2010) and custom Python scripts (https://github.com/thr44pw00d/population-structure). Metagenome/species pairs with an average coverage depth of the reference genome ≥2 (corresponding to ∼20 in the complete metagenome) and a coverage breadth ≥50% were considered potentially useful for population genomic analysis. Only those species that showed a broad geographic range when applying these criteria were used in further analyses. The respective metagenomes were downloaded completely and used for population genomic analyses described below.

### Population genomics

We used POGENOM v0.8.3 (Sjöqvist *et al*., 2021) to determine allele frequencies in the metagenomes and calculate π and F_ST_ values. To generate the therefore necessary vcf files, the *input_pogenom*.*sh* pipeline was run separately for each species, mapping the pre-selected metagenomes against the reference genome. Parameters were set to realize ≥95% identity for the Bowtie2 mapping, a median coverage depth ≥30 and a coverage breadth ≥50% at a mapping quality ≥20 for metagenomes to be included. Metagenome/species pairs that fulfilled these criteria are given in Suppl. Table S1 and population genomic results in Suppl. Table S2. Mapped reads were subsampled to obtain a median coverage depth of 30 for each metagenome. Furthermore, we ran POGENOM with the following tweaks to calculate π based on the actual rather than an estimated number of included loci (newer versions of POGENOM possibly do so by default). Adding the --report-monomorphic flag to the “freebayes_parameters” and ‘QUAL > NA’ as setting for “vcffilter_qual” in the Input POGENOM config file ensured that the pipeline generated a vcf file that contained all loci (including monomorphic loci and without filtering on the probability for loci being polymorphic). The vcf file was subsequently filtered using a custom python script to keep only loci with >99% estimated probability of being mono- or polymorphic, respectively. The *pogenom*.*pl* script was then run twice, first with the “--pi_only” flag. This run was just to get the number of included loci. For the actual run, the “--genome_size” was set to the number of loci determined in the first run. In both runs the filtered vcf file was used, with “--min_count” and “--subsample” set to 15 and “--min_found” to the number of metagenomes. This means that only loci covered at least 15-fold in each metagenome were included, and the coverage at each included locus was subsampled to 15. Linkage disequilibrium was computed from the merged bam files obtained from POGENOM using InStrain v1.6.3 (Olm *et al*., 2021).

### Blast read mapping

For read mapping to reference genomes using blast, each metagenome was subsampled to 1 million reads using seqtk v1.2-r101 (https://github.com/lh3/seqtk). Fastq files were transformed to fasta using Fastx v0.0.14 (https://github.com/agordon/fastx_toolkit), whereby reads containing unknown nucleotides (Ns) were discarded. Read mapping was done as previously described in (Rodriguez-R *et al*., 2021) (settings: -task “blastn” -evalue 0.01 -max_target_seqs 10 -perc_identity 70). Blast results were filtered for a minimum read length of 70 bp and an alignment length ≥90% of the read length using the TabularBlast_ShortRead_Filter.py script (https://github.com/rotheconrad/GoM). Histograms of percent identities of the filtered hits were plotted using the hist function in R v4.2.2 (R Core Team, 2022).

### Exclusion of metagenomes

To obtain the final set of metagenomes used for each species, metagenomes that fulfilled the POGENOM criteria were filtered to exclude those for which a substantial amount of the reads mapped at ≥95% identity that may have originated from a sister species. This was done by removing metagenome/species pairs for which the number of blast hits with 90-95% identity was higher than the number of hits with 95-100% identity. Blast histograms of the final set of metagenome/species pairs are shown in Suppl. Fig. S2 and of the excluded ones in Suppl. Fig. S3, concerning a total of eight metagenomes associated to F. sp., F. ubi., P. ver. and M. uni.

### Calculation of intra-species diversity

Nucleotide diversity (π) and number of polymorphic loci were obtained from POGENOM. ANIr_95_ values were computed from the blast mapping as the average of percent identity values of reads that mapped with ≥95% identity. For the species-wide blast histograms and ANIr_95_ values shown in Fig. 6, the percent identity values from the different metagenomes used for each species were combined.

Estimates of strain diversity were computed as the sum of the Shannon entropies of the allelic frequencies of all the non-correlated polymorphic loci (H_noncorr_) according to the following rationale.

1. Let L_1_ be a polymorphic locus of frequencies [f_i_,f_j_,f_k_,f_l_], sorted from highest to lowest.
  - E.g. L_1_ = [f_A_=0.6, f_C_=0.3, f_T_=0.2, f_G_=0.1].
2. The frequencies of L_1_ in the population can be explained by the presence of four strains with the same frequencies as L_1_ [f_i_,f_j_,f_k_,f_i_], each bearing a different allele in L_1_.
  - E.g. S_1_^L1:A^=0.6, S_2_^L1:C^=0.3, S_3_^L1:T^=0.2, S_4_^L1:G^=0.1).
3. The Shannon strain diversity for that species in that population would thus be H(f_i_,f_j_,f_k_,f_l_), where H is the Shannon entropy of a vector of frequencies.
  - E.g. H(L_1_) = H(0.6, 0.3, 0.2, 0.1) = 1.20
4. Let L_2_ be a second polymorphic locus whose sorted allelic frequencies are similar or highly correlated to those of L_1_.
  - E.g. L_2_ = [f_T_=0.6, f_A_=0.3, f_C_=0.2, f_G_=0.1]).
5. The frequencies of L_1_ and L_2_ in the population can be parsimoniously explained by the same number of strains than the frequencies of L1 alone.
  - E.g. S_1_^L1:A;L2:T^=0.6, S_2_^L1:C;L2:A^=0.3, S_3_^L1:T;L2:G^=0.2, S_4_^L1:G;L2:G^=0.1).
6. Observing several loci with highly correlated sorted allelic frequencies should thus not result in an increased estimated strain diversity.
  - E.g. H_noncorr_(L_1_;L_2_) = H_noncorr_(L_1_)
7. Let L_3_ be a third polymorphic locus whose sorted allelic frequencies [f_i’_,f_j’_,f_k’_,f_l’_] are not correlated to those of L_1_ and L_2_.
  - E.g. L_3_ = [f_G_=0.4, f_T_=0.4, f_A_=0.2, f_C_=0].
8. Two loci with non-correlated sorted allelic frequencies can in most cases not be explained by the same number of strains as a single locus, and thus should result in an increased estimated strain diversity.
  - E.g. H_noncorr_(L_1_;L_2;_L_3_) = H_noncorr_(L_1;_L_2_) + H(L_3_) = 1.20 + 1.05 = 2.25

This is a pragmatic approach meant to provide a fast approximation to intra-species strain diversity without the need for haplotype reconstruction. We benchmarked it using a synthetic dataset, showing that it provides a more accurate representation of strain diversity than π (Suppl. Fig. S5, Suppl. Table S3), which will always increase with the number of polymorphic loci. In this work our threshold for considering two loci as correlated was 1, meaning that only loci with identical sorted allelic frequencies were collapsed, but lower thresholds may be desirable to accommodate noise and sequencing errors, particularly if the coverage depth per locus is higher than the one used in this study.

### Divergence with spatial distance and time

Analyses were done in R v4.2.2. Spatial distances were calculated using the distm function (fun= distGeo) from the geosphere package. Time differences were calculated using the difftime function. To assess correlations of F_ST_ values with spatial distances and time differences, Mantel tests were conducted using the mantel function (method=“spearman”, permutations=10000) from the vegan package. Linear regressions were computed using the stat_smooth function (method=“lm”) from the ggplot2 package. Weighted principal coordinates analyses of F_ST_ values were computed using the wcmdscale function (k=2, eig=TRUE) from the vegan package. Ellipses edging continents and habitats were drawn using the ordiellipse function (scaling=“symmetric”, kind=“ehull”).

### Population dynamics

Allele frequency spectra for the different metagenomes, giving the proportion of each base at each polymorphic locus based on 15-fold coverage, were obtained from the POGENOM results. For clustering alleles in the *P. pan*. and F. sp. time series, Spearman rank correlations between alleles were calculated based on allele frequencies over time using the spearmanr function from the scipy.stats module in Python. All following analyses were done in R. Correlation distances were calculated as 1 – r and visualized in a matrix using the pheatmap function. Correlation distances were clustered using hclust. To infer the number of groups the as.clustrange function from the WeightedCluster package was run for up to 20 groups (ncluster=20) and the number of groups was chosen based on PBC (Point Biserial Correlation). The hierarchically clustered alleles were split into the respective number of groups using the cutree function and the split visualized on the tree using the dendextend package. Dynamics of the groups across the time series were plotted using ggplot2, after calculating the mean allele frequency of all alleles in each group for each metagenome.

## Acknowledgments

We thank Johanna Schmidt for extraction of metagenomic DNA and Alexandra Pitt for help in sampling. Computations were enabled by resources in projects SNIC 2021/5-51, 2021/5-53 and 2021/22-602 provided by the Swedish National Infrastructure for Computing (SNIC) at UPPMAX. This work was supported by the Swedish Research Council (grant 2017-04422), the Swedish research council Formas (grant 2019-02336), the Tiroler Wissenschaftsförderung (project UNI-0404/2370), the Austrian Science Fund (FWF) project 27160-B22, the Research Council of Norway (Project no. 300846), ERA-Net Cofund project BlueBio (grant agreement no. 311913) and a SciLifeLab fellowship to SLG. FP-S was supported by the European Union’s Horizon 2020 research and innovation programme under the Marie Skłodowska-Curie grant agreement No 892961.

## Data Availability

Metagenomic data obtained for this study is available in the NCBI database at https://www.ncbi.nlm.nih.gov under BioProject accession PRJNA965924, SRA accessions SRR24390586-SRR24390649. The data has not yet been released but the metadata can be accessed through the following link: https://dataview.ncbi.nlm.nih.gov/object/PRJNA965924?reviewer=ta1i0hr75on7h2ccng6d1h53g8 Custom scripts used in this study are available on Github at https://github.com/thr44pw00d/population-structure.

## Supplemental Material

**Suppl. Fig. S1**. Detection of each species in a separate map.

**Suppl. Fig. S2**. Blast histograms of all metagenomes per species.

**Suppl. Fig. S3**. Omitted metagenomes due to presence of sister species.

**Suppl. Fig. S4**. ANIr_95_ values with and without metagenomes from the home habitats of the reference genomes.

**Suppl. Fig. S5**. Comparison of H_noncorr_ with different diversity measures in a test dataset.

**Suppl. Fig. S6**. F_ST_ versus spatial distance separated into short (<500 km) and long (≥500 km) distances.

**Suppl. Fig. S7**. Population dynamics in time series metagenomes.

**Suppl. Fig. S8**. Allele frequency histograms for two time series metagenomes.

**Suppl. Fig. S9**. Hierarchical clustering of alleles in two time series based on allele frequency correlations.

**Suppl. Table S1**. Metagenomes used for each species and associated metadata.

**Suppl. Table S2**. Population genomics results for each species and its respective metagenomes.

**Suppl. Table S3**. Dataset used to test H_noncorr_ in comparison to different diversity measures (2 sheets).

**Suppl. Table S4**. Reference genomes used for pre-screening of species in the metagenomes.

**Suppl. Table S5**. Metagenomes generated in this study and associated metadata (2 sheets).

