## Supplementary material for "Geographic population structure and distinct population dynamics of globally abundant freshwater bacteria": Suppl. Fig. S1

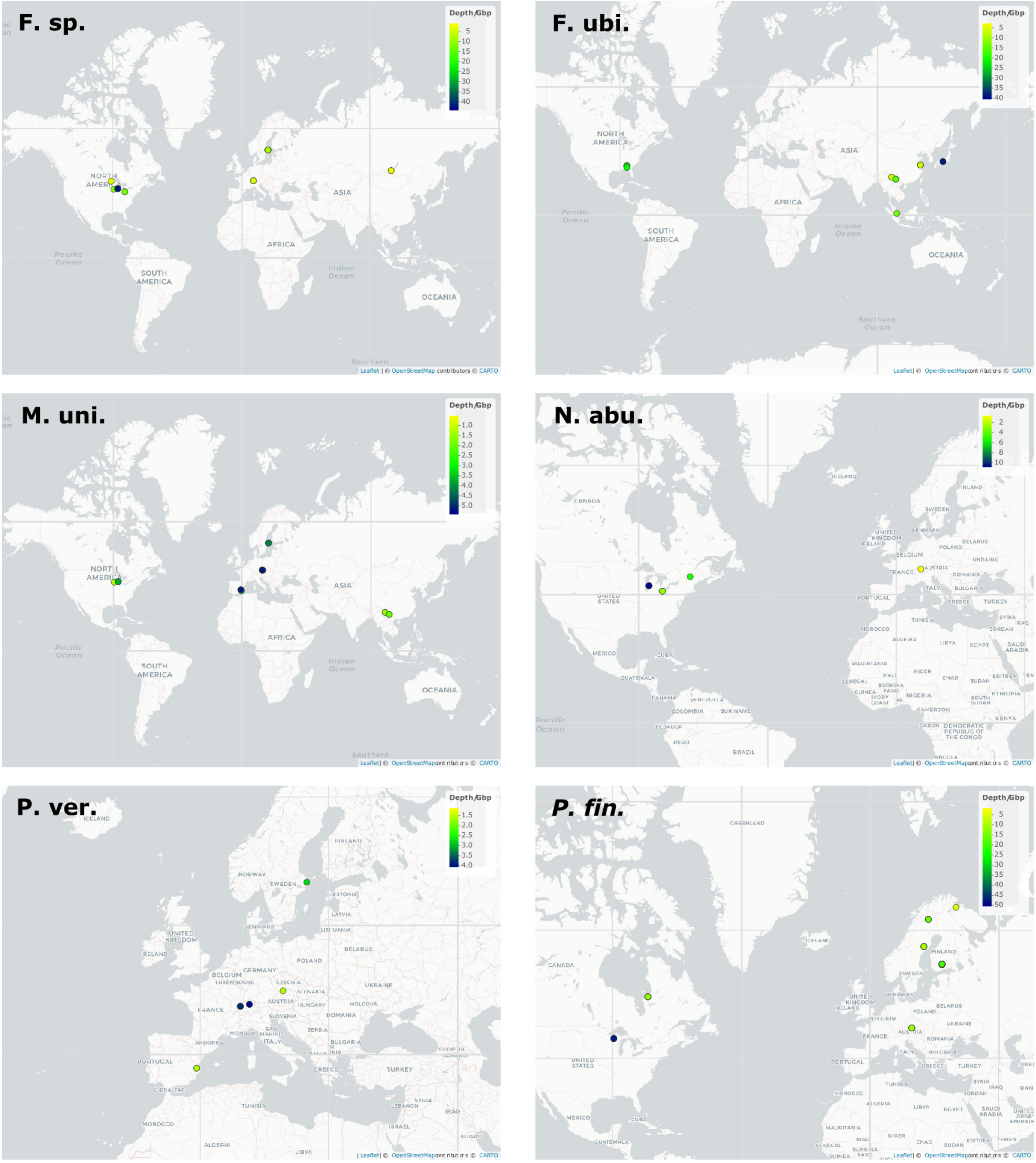


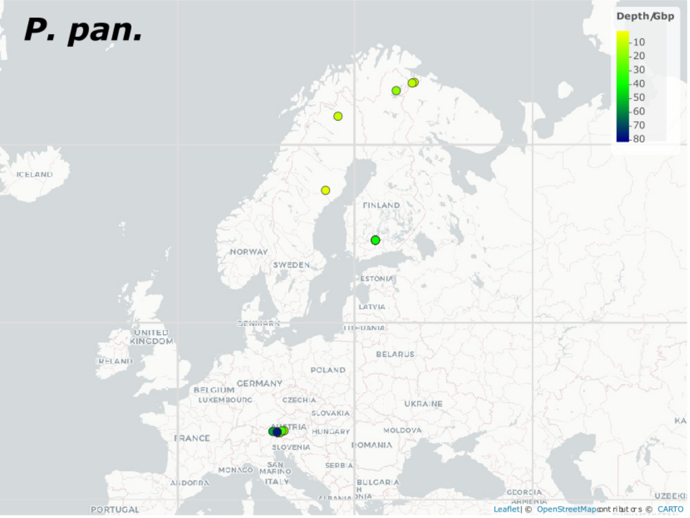


**Suppl. Fig. S1.: Coverage depth of the species in the respective metagenomes.** The sampling sites of the metagenomes used for each species are plotted in separate maps. The color indicates the coverage depth of the reference genome per Gbp of metagenome as obtained from read mapping. Note that not all metagenomes are visible as some dots are overlaid by others.
