## Supplementary material for "Geographic population structure and distinct population dynamics of globally abundant freshwater bacteria": Suppl. Fig. S2

**Suppl. Fig. S2.: Histograms of all metagenomes per species mapped to the respective reference genome by Blast.** The ANIr95 value is given and indicated by a blue dashed line for each histogram. The histograms are sorted by increasing spatial distance between the sampling site of the metagenome and the home habitat of the reference genome. The respective distances are given in km next to the metagenome name on top of each histogram.

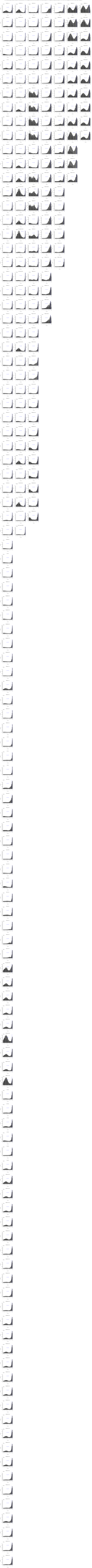
