## Supplementary material for "Geographic population structure and distinct population dynamics of globally abundant freshwater bacteria": Suppl. Fig. S3

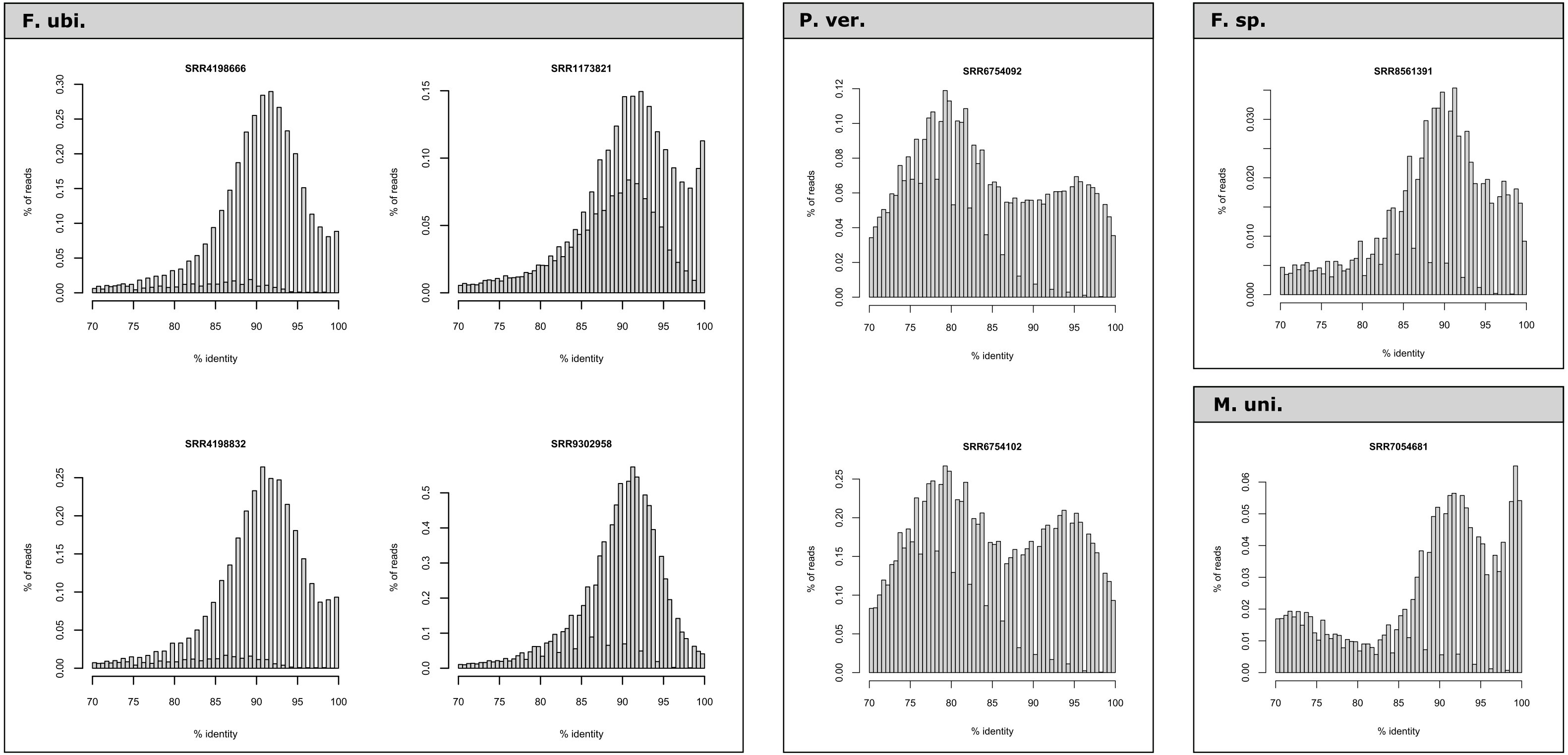


**Suppl. Fig. S3.: Omitted metagenomes due to presence of sister species.** These eight metagenomes were excluded from analyses of the respective species as the number of reads mapped with 90-95% identity was higher than the number of reads with >95% identity, i.e. a substantial fraction of the latter reads may have originated from a sister species rather than the species of interest.
