## Supplementary material for "Geographic population structure and distinct population dynamics of globally abundant freshwater bacteria": Suppl. Fig. S4

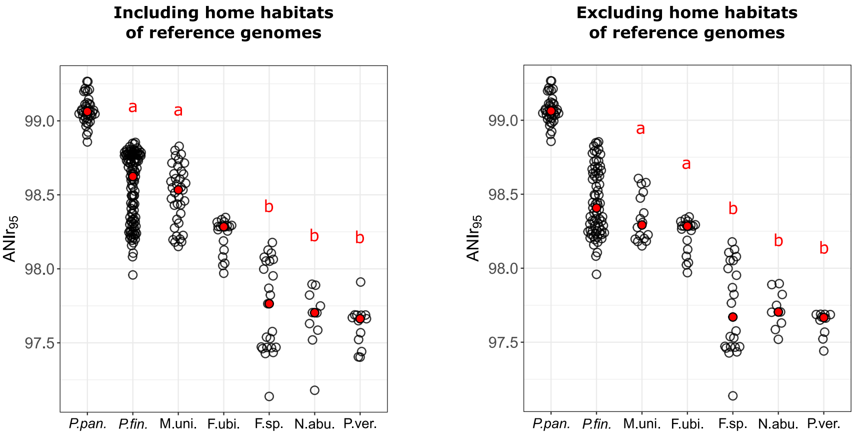


**Suppl. Fig. S4.: ANIr$_{95}$ values for the seven species when metagenomes sampled from the habitat where the reference genome was obtained from are included (left) and excluded (right).** Each dot refers to one metagenome. Red dots show the medians. Pairs within each plot that are not significantly different ($p > 0.05$) according to Wilcoxon-Mann-Whitmey Rank Sum test are marked with a common letter.
