## Supplementary material for "Geographic population structure and distinct population dynamics of globally abundant freshwater bacteria": Suppl. Fig. S5

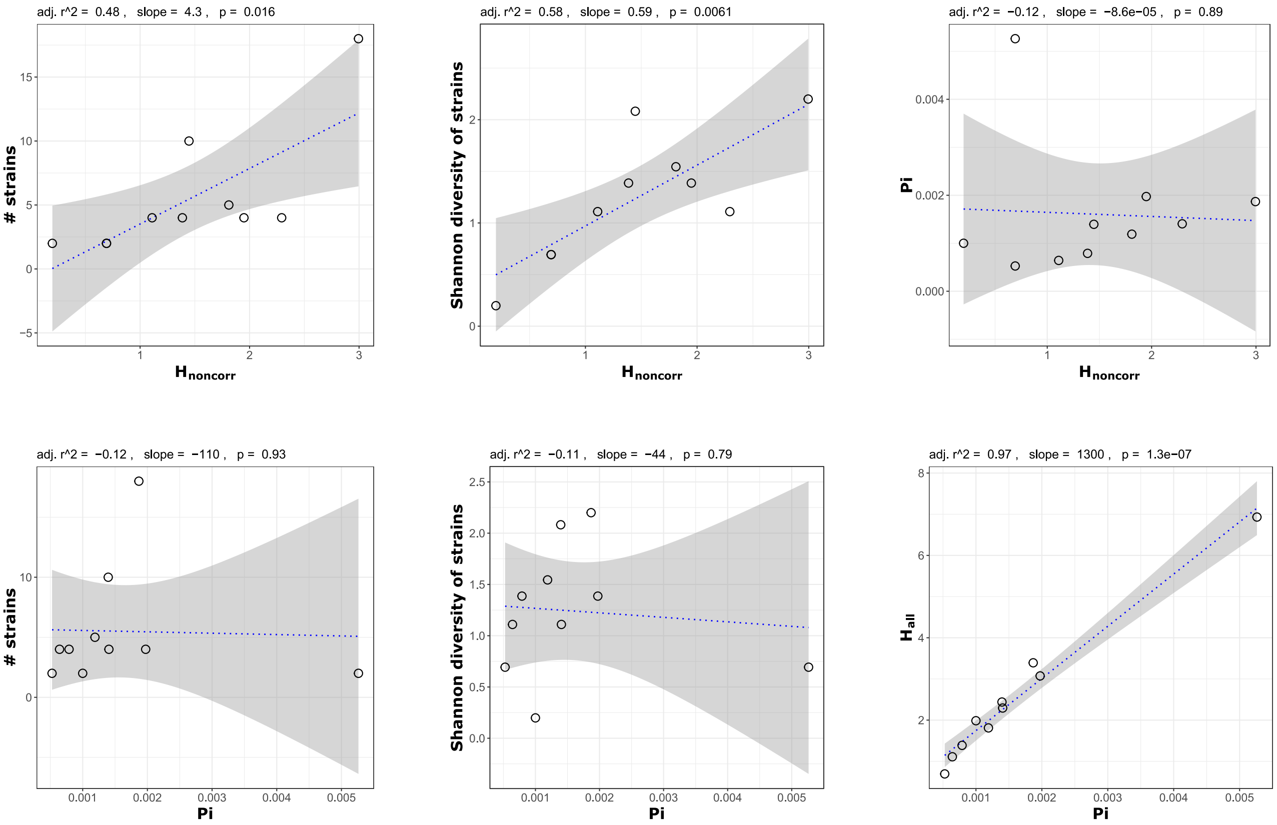


**Suppl. Fig. S5.: Comparison of different diversity measures in test dataset.** The Shannon entropy of polymorphic loci with non-correlated allele frequencies ($H_{noncorr}$) correlates better with the number of strains than Pi. The latter diversity measure is strongly correlated with the Shannon entropy of all polymorphic loci ($H_{all}$), i.e. it increases with the number of polymorphic loci regardless of correlation among them.
