## Supplementary material for "Geographic population structure and distinct population dynamics of globally abundant freshwater bacteria": Suppl. Fig. S6

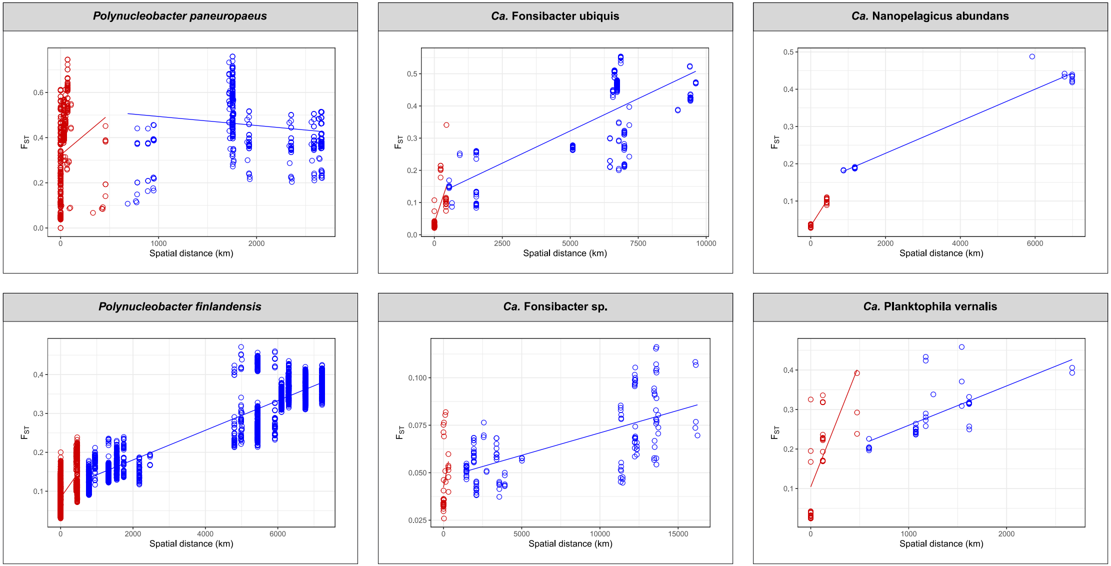


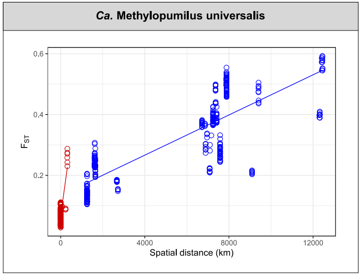


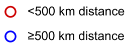


**Suppl. Fig. S6.: Population differentiation ($F_{ST}$) versus spatial distance between habitats separated into short (<500 km) and long (≥500 km) distances.** The increase of $F_{ST}$ with spatial distance is stronger for short distances. Short-range dispersal is probably more affected by direct dispersal events between habitats. As the magnitude of such dispersal events is expected to decrease quadratically with distance it is not surprising that the rate of $F_{ST}$ increase is larger at shorter distances. However, a more linear increase of $F_{ST}$ with spatial distance is apparent over long distances. This might be explained by incremental dispersal events between nearby habitats being more effectual than direct dispersal over long distances (stepping-stone model).
