## Supplementary material for "Geographic population structure and distinct population dynamics of globally abundant freshwater bacteria": Suppl. Fig. S7

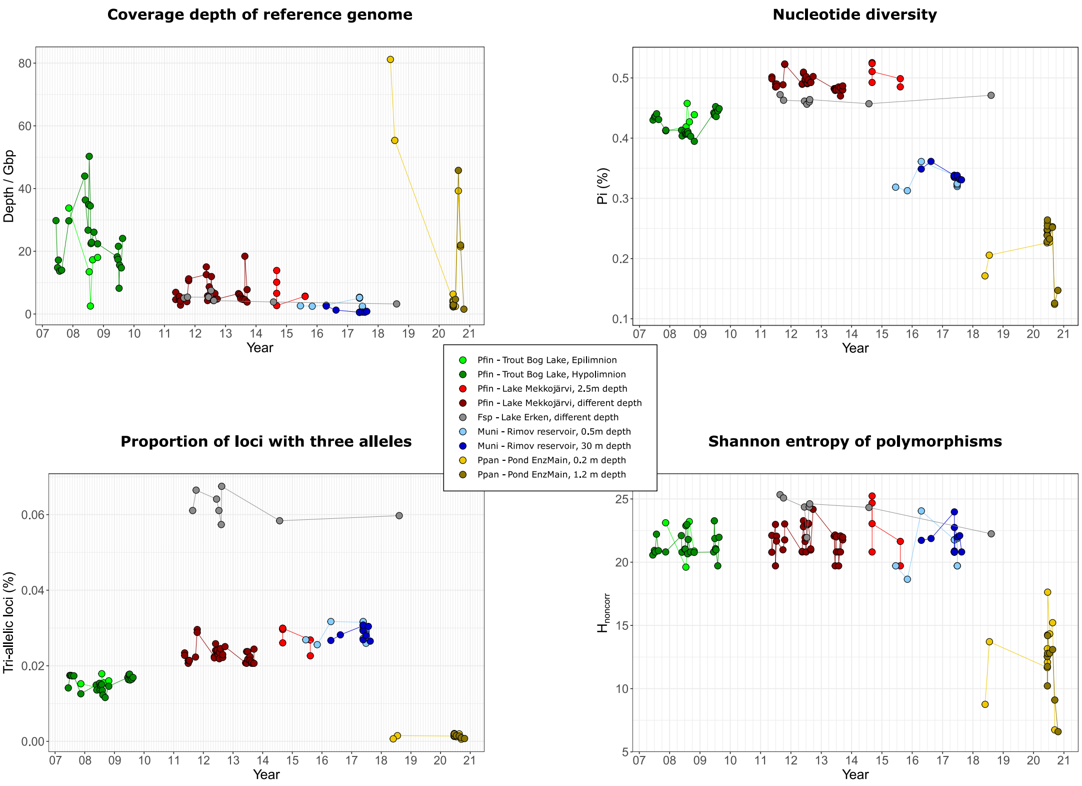


**Suppl. Fig. S7.: Population dynamics in time series metagenomes.** Relative abundance inferred from read mapping (A) and different diversity measures (B, C, D) in time series for four species in five different habitats. Data points for metagenomes from the same habitat but sampled from different depths are depicted by similar color but different brightness.
