## Supplementary material for "Geographic population structure and distinct population dynamics of globally abundant freshwater bacteria": Suppl. Fig. S8

### *Polynucleobacter paneuropaeus* - Pond EnzMain

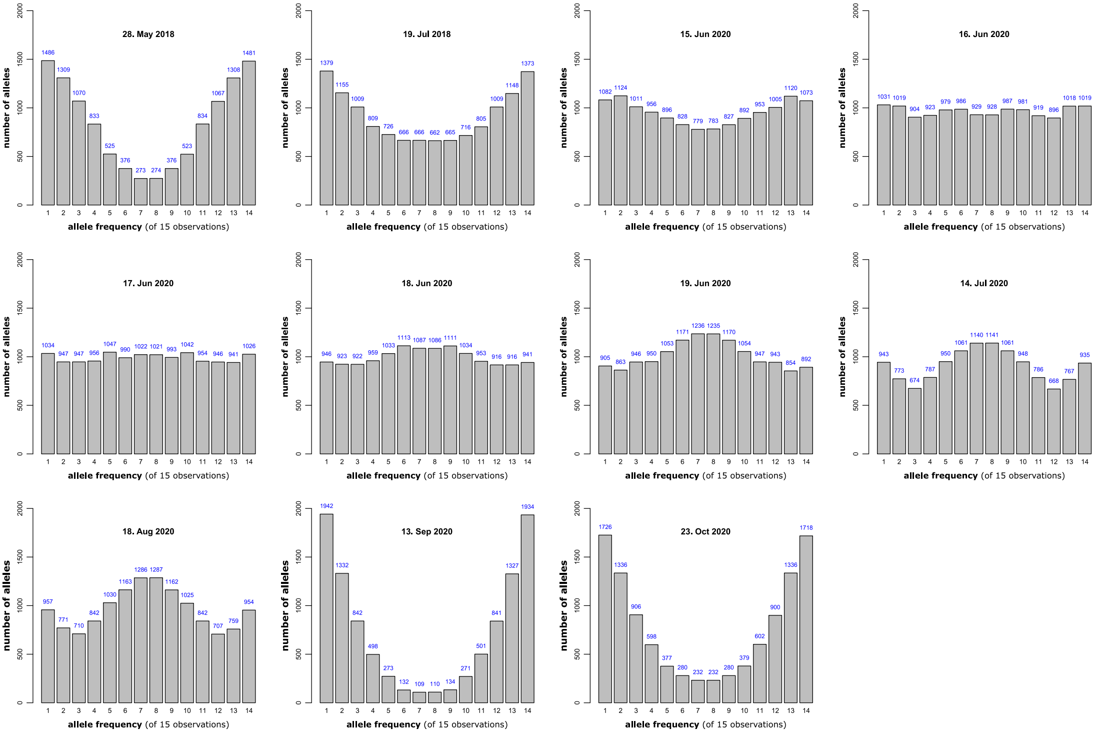


### *Ca.* Fonsibacter sp. - Lake Erken

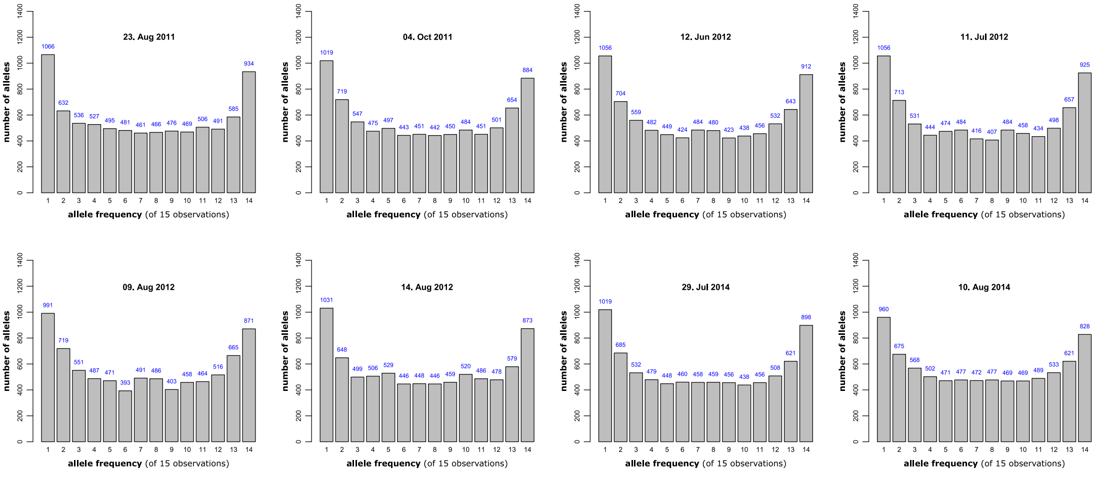


**Suppl. Fig. S8.: Allele frequency histograms from time series metagenomes for two species in two different habitats.** The *Polynucleobacter paneuropaeus* population in Pond EnzMain showed strong variation among different timepoints. Times where one genotype seemed to dominate the population (May and July 2018, September and October 2019) were intermitted by times when at least two different genotypes were dominant (June, July and August 2020). The *Ca.* Fonsibacter sp. population showed similar allele frequencies at all timpoints. Possibly, multiple genotypes coexisted steadily over multiple years.
