## Supplementary material for "Geographic population structure and distinct population dynamics of globally abundant freshwater bacteria": Suppl. Fig. S9

***Polynucleobacter paneuropaeus* - Pond EnzMain**

Split into 9 groups based on Point Biserial Correlation cluster quality

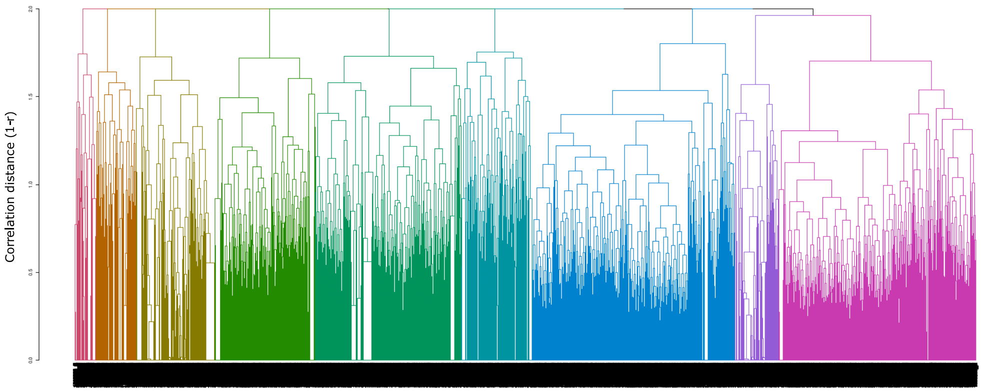


***Ca.* Fonsibacter sp. - Lake Erken**

Split into 11 groups based on Point Biserial Correlation cluster quality

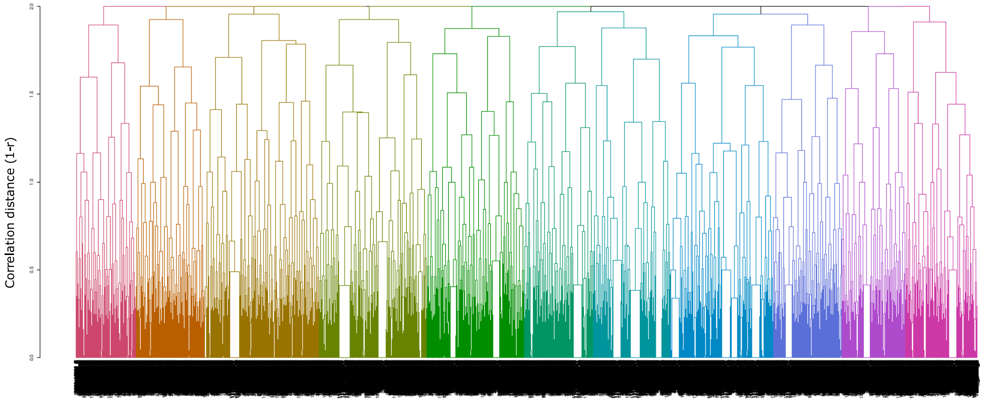


**Suppl. Fig. S9.: Hierarchical clustering of alleles based on correlation of allele frequencies across time series of two species in two different habitats.** Alleles of the *Polynucleobacter paneuropaeus* population were clustered into nine and those of the *Ca.* Fonsibacter sp. population into eleven groups based on Point Biserial Correlation cluster quality. It is apparent that the groups in *Polynucleobacter paneuropaeus* are more distinct than those in Ca. Fonsibacter sp., pointing at more distinguished genotypes in the former and a more continuous diversity structure in the latter population.
